# NEK7 phosphorylation of cortactin modulates the migratory capacity of cells expressing EML4-ALK V3

**DOI:** 10.1101/2025.08.30.672807

**Authors:** Emily L. Richardson, Axel Knebel, Kees R. Straatman, Robert Gourlay, Douglas Lamont, Tara Hardy, Robert E. Turnbull, Susan W. Robinson, Laura O’Regan, Richard Bayliss, Andrew M. Fry

## Abstract

EML4-ALK is a common oncogenic driver of non-small cell lung cancer. Distinct EML4-ALK variants cause different rates of disease progression, with patients expressing variant 3 (V3) exhibiting accelerated metastasis. Cells expressing EML4-ALK V3 develop a mesenchymal-like morphology and enhanced migration that is dependent on the NEK9 and NEK7 kinases. However, downstream substrates of these kinases relevant to these phenotypes are largely unknown. Here, we show that the actin-binding protein cortactin is phosphorylated by NEK7 within the F-actin-binding region (ABR) and that depletion of cortactin abrogates the morphological and migration phenotypes induced by EML4-ALK V3. Expression of constitutively active mutants of NEK9 or NEK7 causes similar cortactin-dependent morphological and migration changes. Cortactin co-localises with NEK7 and EML4-ALK V3 at branched filopodia-like extensions that are also generated upon expression of a cortactin protein with phospho-mimetic mutations in the ABR. In contrast, phospho-null mutations dissociate cortactin from F-actin. We propose that EML4-ALK V3 alters cell morphology and promotes directed cell migration by modulating the actin cytoskeleton via NEK7-mediated phosphorylation of cortactin within its ABR.

**SUMMARY STATEMENT:** NEK7 phosphorylates key residues of cortactin in its actin-binding repeat region to modulate the actin architecture and migration of cells driven by the EML4-ALK V3 – NEK9 – NEK7 pathway.

## INTRODUCTION

Effective cellular migration requires exquisite control of numerous effector proteins and cytoskeletal elements, together acting in concert to direct cells towards stimuli. In cancer, migration is required for cancer cells to migrate out of the primary tumour in the process of metastasis – the leading cause of cancer deaths. The deadliest cancers are those that metastasise quickly and effectively, often through manipulating normal pathways to act in an aberrant manner.

Non-small cell lung cancer (NSCLC) constitutes approximately 85% of the 2.2 million lung cancer diagnoses worldwide ^[1]^. Importantly, at point of diagnosis close to 50% of patients present with distant metastases, which correlates with under 30% average 5-year survival rate of NSCLC patients ^[2–4]^. Approximately 5% of NSCLCs are driven by a fusion of two unrelated proteins, echinoderm microtubule associated-like protein 4 (EML4) and anaplastic lymphoma kinase (ALK) ^[5]^. This fusion can arise through distinct breakpoints within the *EML4* gene, resulting in the generation of different EML4-ALK variants with more than fifteen identified to date ^[6]^.

EML4-ALK variant 1 (V1) and variant 3 (V3) are the most common and represent more than 80% of cases ^[7]^. Importantly, these two variants lead to very different clinical outcomes and response to targeted therapies, with patients expressing EML4-ALK V3 exhibiting more aggressive metastatic tumours that respond much less well to ALK inhibitors than patients expressing EML4-ALK V1^[8,9]^. Coinciding with these different clinical outcomes, a number of differences in the biological properties of EML4-ALK V1 and V3 have been reported, including their protein stability and subsequent microtubule binding ability, formation of signalling liquid-liquid phase separated droplets, and interaction with other proteins^[10–13]^.

Corresponding to this, the expression of EML4-ALK V3, but not V1, leads to development of a mesenchymal-like cell morphology with production of long cytoplasmic protrusions, as well as an increased rate of cell migration and invasion ^[12]^. These phenotypes are dependent upon the NEK9 and NEK7 kinases with which EML4-ALK V3 but not V1 can interact ^[12]^. This interaction occurs via the N-terminal region of EML4 and leads to recruitment of NEK9 and NEK7 to microtubules in cells expressing EML4-ALK V3 ^[12,14]^. Human cells express eleven NEK kinases that share approximately 40-50% sequence conservation within their catalytic domains^[15]^. NEK7 is most closely related to NEK6, with both kinases consisting of little more than a catalytic domain that shares 85% identity. Both NEK6 and NEK7 have roles within mitosis regulating mitotic spindle assembly and cytokinesis ^[16–19]^. The kinase activities of NEK6 and NEK7 are stimulated by NEK9, which is downstream of CDK1 and PLK1^[15,17,20–22]^.

Intriguingly, through an *in vitro* kinase-substrate screening assay we had previously used to identify the heat shock protein HSP72 as a NEK6 substrate (O’Regan et al. 2015) ^[18]^, we also discovered cortactin as a potential substrate of NEK6. Cortactin is an actin binding protein that is well characterised as a cellular marker of advanced disease in a diverse range of cancers ^[23,24]^. It has also been demonstrated to have key roles in the lung, particularly for maintenance of barrier integrity and migration of cells in response to damage ^[25,26]^.

Cortactin is a class 2 nucleation-promoting factor (NPF) that binds to F-actin and the Arp2/3 complex, enabling the Arp2/3 complex to catalyse the formation of branched F-actin through promoting actin nucleation and elongation. The binding of cortactin to F-actin occurs through an F-actin binding region (ABR) that stabilises F-actin filaments as well as promoting the formation of a branched actin network via its N-terminal interaction with Arp3 ^[27,28]^. These actions of cortactin have important roles at distinct migratory and invasive structures as well as in cell division ^[29–32]^.

The cortactin ABR consists of six 37 amino acid repeats with an incomplete seventh repeat of 20 amino acids ^[33]^. It has been shown that the fourth repeat is the most important for F-actin binding ^[34]^. In particular, a serine residue within the fourth repeat, Ser209, has been reported to have a specific role in actin binding and stabilisation ^[35]^. Here, we show that members of the NEK kinase family, NEK6 and NEK7, phosphorylate cortactin within the ABR, including at Ser209, thus having the potential to alter actin binding and architecture. Furthermore, we establish that cortactin is required for the effective migration of cells expressing EML4-ALK V3, with our results supporting a model in which this activity is mediated via interaction with and phosphorylation by NEK7.

## RESULTS

### NEK6 and NEK7 phosphorylate cortactin with varying affinities

To identify NEK6 substrates, we had previously undertaken a KinasE Substrate Tracking and Elucidation (KESTREL) screen in which fractionated HEK293 cell lysates were incubated with purified NEK6 kinase in the presence of Mn^2+^ and [γ^32^P]-ATP to detect proteins phosphorylated with high specific activity. Following several rounds of fractionation, proteins that exhibited substantial phosphorylation were cut out of an SDS-PAGE gel and identified by mass spectrometry. This led to discovery of HSP72, as well as β-tubulin, as substrates of NEK6 ^[18]^. However, of all the proteins identified in this assay, the substrate with the highest rate of initial phosphorylation was the actin-associated protein, cortactin (Fig. S1A & B).

To determine whether cortactin was limited to NEK6 as a substrate, we investigated whether NEK7, a close relative of NEK6 with >85% homology within their catalytic domains, was also able to phosphorylate cortactin. Using *in vitro* kinase assays with cortactin protein expressed and purified from bacteria, it was determined that not only did NEK7 phosphorylate cortactin, but it did so more rapidly and to a greater extent than NEK6 (Figure 1A & B). The rates of kinase activity were analysed and the V_max_ of NEK7 determined to be >10-fold higher than that of NEK6 (Figure 1C). Additional kinase assays were completed to further characterise the results against a known substrate of NEK6, HSP72 ^[18]^. Indeed, whilst HSP72 was a better substrate for NEK6 than NEK7, as previously demonstrated, cortactin was phosphorylated more strongly by NEK7 (Fig. S1C).

**Figure 1.**
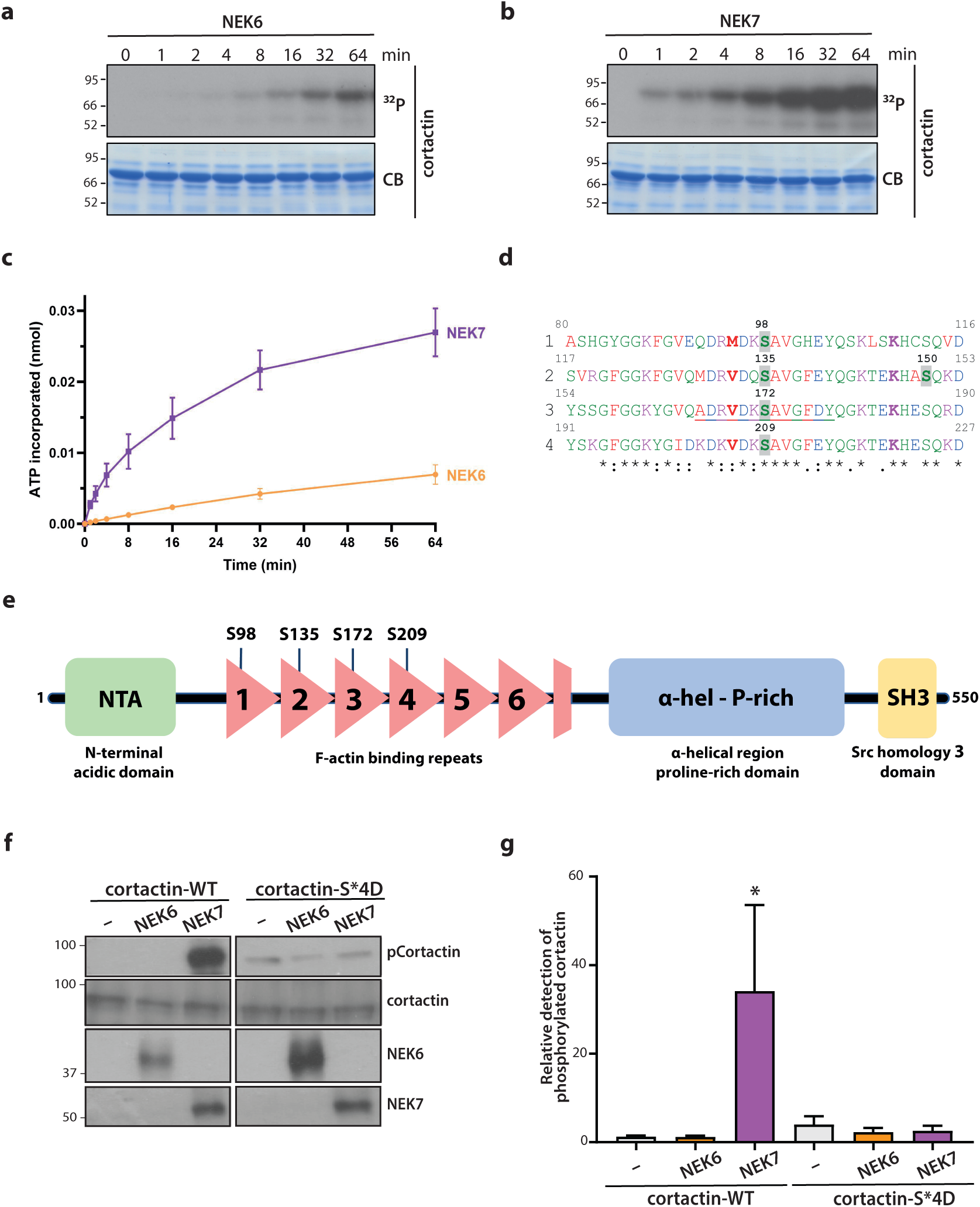
NEK6 and NEK7 phosphorylate the actin-branching protein, cortactin in vitro. **A & B.** Recombinant human cortactin A was incubated with purified His-tagged NEK6 (A) or NEK7 (B) protein in the presence of 10 mM MgCl_2_ and 0.1 mM γ^32^P-ATP for the indicated times before reactions were quenched with SDS-loading buffer and boiled for 5 mins. Samples were separated on 4%-20% Tris-Glycine SDS-gels and stained with Instant Coomassie Blue (CB) and exposed to X-ray film (^32^P). **C.** Cortactin proteins were excised from the gel and radioactive decay measured in a Scintillation Counter. The amount (nmol) of ATP incorporated was plotted against time. Vmax was determined to be 0.21 pmol min^-1^ for NEK6 and 2.65 pmol min^-1^ for NEK7. **D.** Multiple sequence alignment of the first four F-actin binding repeats as determined using Clustal Omega (EMBL-EBI) is presented and residue conservation ranked as indicated: [*] fully conserved; [:], strong conservation; [.], weak conservation. Residue characteristics are indicated using the following colours: small hydrophobic residue (red), acidic residue (blue), basic residue (magenta), hydroxyl/sulfhydryl/amine residue (green). Residues phosphorylated by NEK7 are highlighted. The hydrophobic residue at -3 typical of the NEK6 and NEK7 consensus sequence is marked in bold. The underlined residues indicate the peptide sequence used for generation of the pCortactin antibody. **E.** Schematic cartoon of the cortactin protein indicating the N-terminal acidic domain (NTA), the 6.5 F-actin binding repeats, α-helical region and proline-rich domain (α-hel - P-rich) and Src homology 3 (SH3) domain. The four conserved residues phosphorylated by NEK7 in the first four actin binding repeats are indicated. **F.** Recombinant NEK6 and NEK7 kinases were pre-incubated in kinase buffer for 1 hr before being subsequently incubated with purified cortactin-WT or -S*4D proteins for 6 hr. Samples were separated by SDS-PAGE and analysed by Western blot using the pSer172 phospho-antibody (pCortactin), as well as antibodies against cortactin, NEK6 and NEK7. **G.** Quantification of data from F showing the intensity of phosphorylated cortactin (WT or S*4D) relative to total cortactin as measured by ImageJ following incubation with or without the kinase indicated. Graph shows means and standard error (n=3). M. wts (kDa) in A and F are indicated on the left.

Phosphosite mass spectrometry analysis of the purified protein revealed multiple sites of cortactin phosphorylation by NEK6 and NEK7. Interestingly, there was a significant overlap in these sites, although a higher number of sites were phosphorylated by NEK7 (22 sites) than NEK6 (10 sites), consistent with more extensive phosphorylation of cortactin by NEK7 (Fig. S1D & E). Of note, phosphorylation site mapping by mass spectrometry does not provide information about the stoichiometry of phosphorylation on each site, and we assume that many of the identified sites are sub-stoichiometrically phosphorylated.

In terms of protein localisation, NEK6 phosphorylation sites resided within the ABR and the proline rich region, whilst NEK7 sites were distributed across the whole cortactin protein, including the N-terminal acidic domain, ABR and proline-rich region, but excluding the SH3 domain. Seven residues were phosphorylated by both kinases, with four residing in the ABR (Ser135, Ser172, Ser261, Ser277) and 3 in the proline rich region (Thr401, Ser405, Ser432). Phosphorylation of Ser405 by ERK1/2 has previously been reported to promote N-WASP binding and activation ^[31,36]^.

Intriguingly, four of the phosphorylation sites catalysed by NEK7 (Ser98, Ser135, Ser172 and Ser209) are located in a conserved sequence in the first four repeats of the ABR (Fig. 1D & E) ^[35]^. These sit in a preferred consensus phosphorylation sequence for NEK6 and NEK7 with a hydrophobic residue at -3 to the phosphoserine^37,38^. These phosphorylated serine residues also sit 11 residues upstream of a conserved lysine residue that undergoes acetylation to regulate actin binding ^[39,40]^. Interestingly, equivalent serine residues are not found at this position within the fifth, sixth or final half repeat of the ABR.

To further investigate these particular phosphorylation events, a synthetic peptide representing the sequence encompassing the phosphorylated Ser172 residue was used to generate a phospho-specific cortactin (pCortactin) antibody. It is worth noting that the peptide sequence used for immunization is, as previously indicated, highly conserved (albeit not identical) with the sequence around Ser98, Ser135 and Ser209 (Fig. 1D). Hence, the antibody generated may detect phosphorylation at any or all four of the serines in these F-actin binding repeats. Site-directed mutagenesis was used to create a phospho-mimetic S*4D mutant cortactin protein in which all four of these serine residues (Ser98, Ser135, Ser172, Ser209) were mutated to aspartic acid. This alongside a wild type (WT) cortactin protein was expressed and purified from bacteria.

The S*4D and WT cortactin proteins were incubated with NEK6 or NEK7 in kinase assays and then analysed by Western blot with the pCortactin antibody. The phospho-mimetic cortactin-S*4D protein, but not the wild-type protein, was weakly detected by the pCortactin antibody in both the presence and absence of both kinases, suggesting that the phospho-mimetic protein mimics the phosphorylated wild-type protein sufficiently well to be detected by the antibody (Fig. 1F). After incubation with NEK7, a strong band was detected for WT cortactin with the pCortactin antibody. Indeed, there was a striking difference upon incubation with NEK6 and NEK7, with around 20-fold higher detection after incubation with NEK7 than NEK6 (Fig. 1F & G). These results can be explained either by NEK7 phosphorylating Ser172 more efficiently than NEK6, or by the fact that this antibody may also detect the related sites in the ABR phosphorylated by NEK7. Taken together, these biochemical data point to a novel and intriguing association between NEK7 and cortactin.

### Cortactin is required for altered morphology and enhanced migration in cells expressing activated NEK9 or NEK7

Having established that cortactin can be phosphorylated by NEK7, we next asked whether it might be involved in the cellular phenotypes induced by expression of constitutively activated versions of NEK7 or its upstream activator NEK9. For this, we examined the consequences of its depletion by confocal microscopy in stable U2OS cell lines that had been previously engineered to express constitutively active mutants of NEK7 (aNEK7; Y97A) or NEK9 (aNEK9; ΔRCC1) ^[20,21]^. Expression of these activated kinases were previously shown to cause interphase cells to form extended cytoplasmic protrusions and exhibit an enhanced rate of migration ^[12]^. We were able to recapitulate these findings and determined that the expression of the active form of these kinases induced a mesenchymal morphology with altered actin microarchitecture, with cortactin localised throughout the extended cytoplasmic protrusions (Fig. S2A-D).

Two independent siRNAs were used to probe the function of cortactin (siCTTNa and siCTTNb). siCTTNb was shown to induce more effective depletion of cortactin than siCTTNa (Figure 2A) and this difference was utilised throughout the study to demonstrate partial and complete depletion of cortactin and the resulting phenotypes. Depletion of cortactin using two siRNAs led to a abrogation of cytoplasmic protrusions in aNEK9 cells (Fig. 2B-C). The depletion of cortactin in aNEK7 cells led to larger, flatter cells, which also lacked cytoplasmic protrusions as demonstrated by a decreased length-to-width ratio (Fig. 2D-F). In both cell types, cortactin depletion led to significant increases in stress fibres, with quantification demonstrating an increase in length and decrease in curvature of F-actin (Fig. S3A-D).

**Figure 2.**
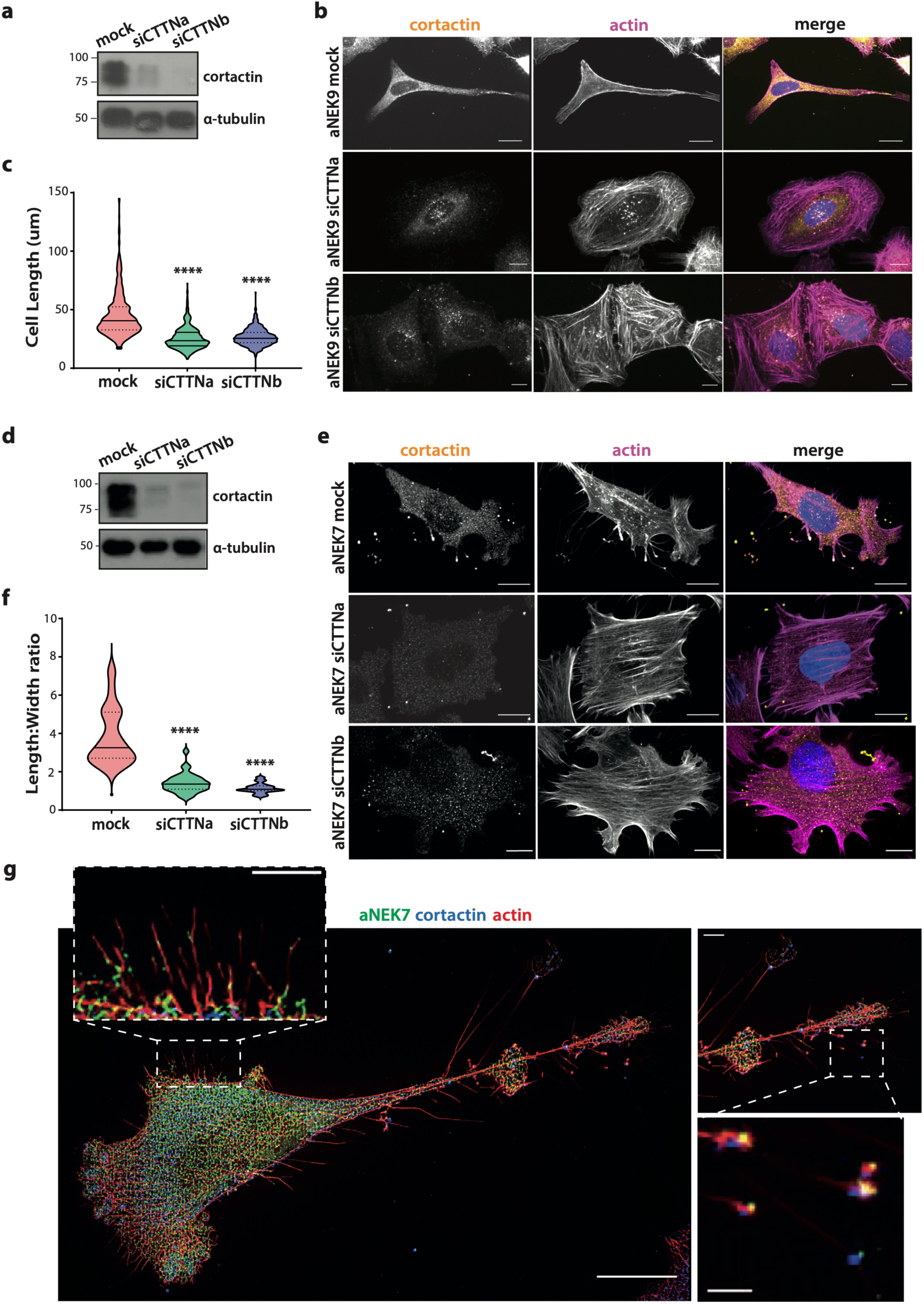
Cortactin is required for the mesenchymal morphology displayed by cells expressing activated NEK9 or NEK7. **A.** U2OS:myc-NEK9-ΔRCC1 cells were induced with doxycycline for 72 hr to express activated NEK9 (aNEK9) and simultaneously were either mock depleted or depleted of cortactin using siCTTNa or siCTTNb. Cell lysates were analysed by Western blot with antibodies against cortactin and α-tubulin. **B.** Cells treated as in A were fixed and stained with cortactin (yellow) antibodies and TRITC-phalloidin (actin, magenta). Merge images include DNA stained with Hoeschst 33258 (blue). **C.** The length of aNEK9 cells treated as in A was measured as described in Materials and Methods. Solid line represents the mean and dotted lines the quartiles (n=3). **D.** HeLa:YFP-NEK7-Y97A cells were induced to express activated NEK7 (aNEK7) and depleted and analysed as described in A. M. wts (kDa) are indicated on the left in A and D. **E.** Cells treated as in D were fixed and stained as in B. Scale bars in B and E, 10 μm. **F.** The length and width of aNEK7 cells depleted of cortactin were measured as described in Materials and Methods and plotted as a ratio. Solid line represents the mean and dotted lines the quartiles (n=3). **G.** HeLa:YFP-NEK7-Y97A cells were induced to express aNEK7 for 72 hr before being fixed and stained with GFP (aNEK7, green) and cortactin (blue) antibodies, and TRITC-phalloidin (actin, red). 50 images were taken and reconstructed using NanoJ-SRRF. Magnified views of the areas indicated in boxes are shown. Scale bars, 20 μm (central image), 5 μm (top left and upper right zoom) and 2 μm (bottom right zoom). ****, *p* < 0.0001 by unpaired Student’s T-test.

Intriguingly, cells expressing aNEK7 were observed to form many fine filopodia-like extensions (FLEs) on the surface of the plasma membrane (Fig. 2E). Super-resolution radial fluctuation (SRRF) imaging of these cells revealed that these were present on both lamellipodia-like surfaces and cytoplasmic protrusions, where they were often branched with multiple tips and could reach >10 µm in length (Fig. 2G). Live cell phase contrast imaging also allowed detection of similar structures suggesting that they were not artefacts of fixation (Movies S1, 2), while fluorescence staining revealed that, although the body of these extensions contained mainly actin, the tips and branch points had detectable foci of both activated NEK7 and cortactin as well as phosphorylated cortactin (Fig. 2G, Fig. S4A). Quantification confirmed significantly increased colocalization of activated NEK7 with cortactin and pCortactin in FLEs compared to the cell body (Fig. S4B-C).

Depletion of cortactin from cells expressing aNEK9 or aNEK7 also significantly reduced their rate of migration as determined by three independent measures. First, using cells expressing aNEK9, cortactin depletion substantially interfered with (i) the rate of wound closure in a scratch wound assay that measures collective migration (Fig. 3A-B); (ii) distance travelled when tracking single cells in a sparsely plated population by time-lapse imaging (Fig. 3C); and (iii) directional migration as measured by Real Time Cell Analysis (RTCA), a modified Boyden chamber assay that detects cells crossing a membrane towards a chemo-attractant (Fig. 3D). A similar set of results was obtained in each of these three assays upon depletion of cortactin in cells expressing aNEK7 (Fig. 3E-H). These data provide persuasive evidence that cortactin is required not only for the altered cell morphology but also the accelerated migration induced by expression of constitutively active mutants of NEK9 or NEK7.

**Figure 3.**
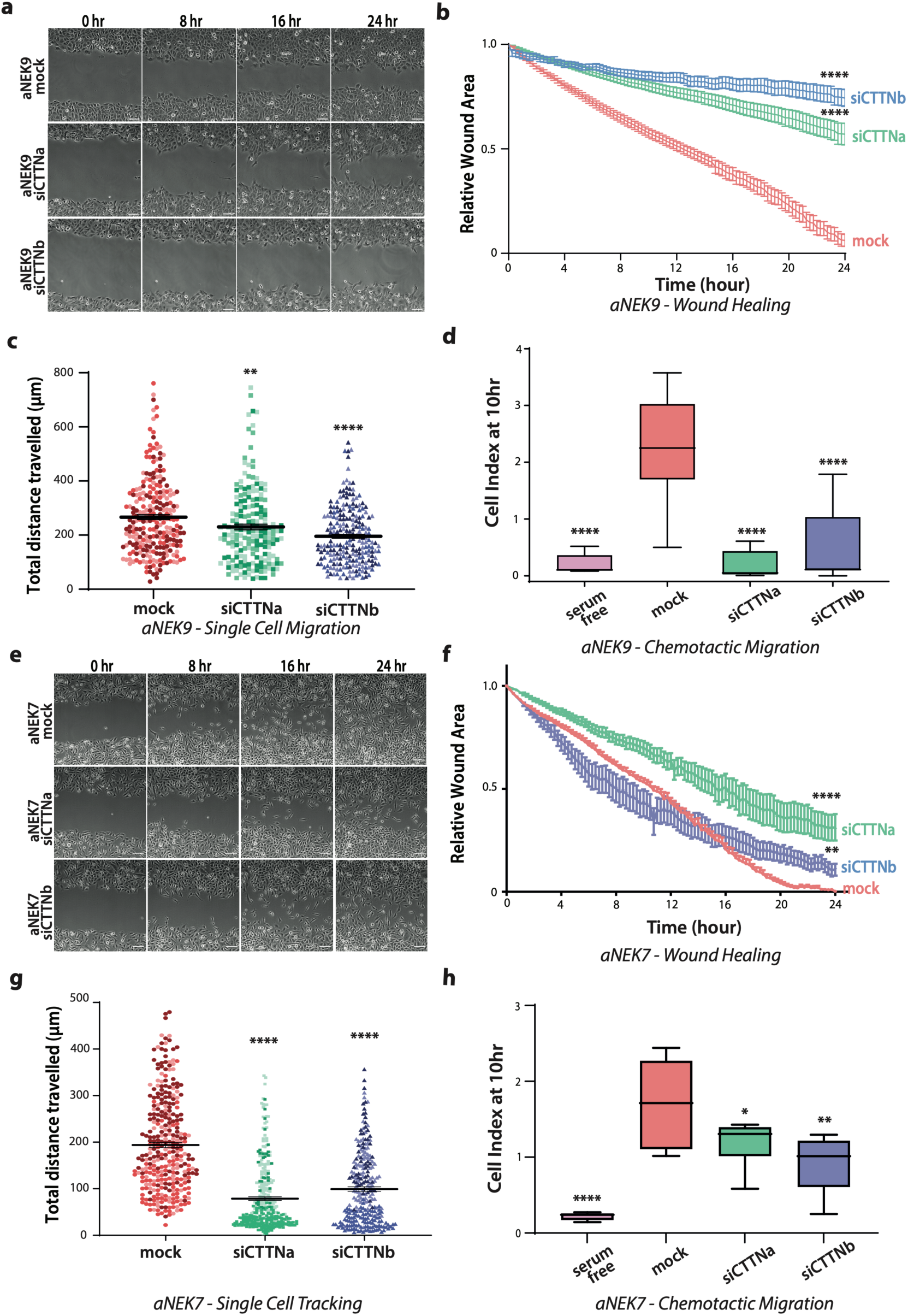
Cortactin is required for migration of cells expressing activated NEK9 or NEK7. **A-D**. U2OS:myc-NEK9-ΔRCC1 cells were induced to express activated NEK9 (aNEK9) and either mock-depleted or depleted of proteins using siCTTNa or siCTTNb siRNA oligonucleotides for 72 hr. **E-H.** HeLa:YFP-NEK7-Y97A cells were induced to express activated NEK7 (aNEK7) and either mock-depleted or depleted of cortactin using siCTTNa and siCTTNb siRNA oligonucleotides for 72 hr. For wound healing experiments (A, E), treated cells were plated in a confluent layer before a single scratch wound was made through the centre of the well. Movement of cells was measured by time-lapse microscopy over 24 hr. Representative images at time points indicated are shown in A and E for aNEK9 and aNEK7 cells, respectively. Scale bar, 100 μm. (B, F) The relative wound area (area devoid of cells) was measured over 24 hr, with mean and SEM indicated. The wound areas at 24 hours were analysed for significant differences (n=3, with 6 replicates per experiment). For single cell migration experiments (C, G), treated cells were plated on collagen-coated wells and distance travelled by individual cells measured over 12 hr by time-lapse microscopy. Each shade of colour within plots represents a biological repeat (n=3, 6 replicates per experiment). The mean and SEM are indicated. For chemotactic migration experiments (D, H), treated cells were placed in the upper chamber of a CIM-plate 16 without serum. The lower chamber was filled with media with serum as a chemo-attractant and the movement of cells across the chamber measured over 48 hr. The cell index at the 10 hr time-point is shown. The mean is indicated (n=3-5 with 2 repeats per experiment). Migration of mock-depleted cells with serum-free media in the lower chamber was also measured. **, *p*<0.01, **, *p<*0.001, ****, *p*<0.0001 by one-way ANOVA.

### Cortactin is required for altered morphology and increased migration in cells expressing EML4-ALK V3

We next investigated whether cortactin is also required for the mesenchymal-like morphology and accelerated migration in U2OS cells expressing EML4-ALK V3, as these phenotypes were previously shown to be dependent on NEK9 and NEK7 activity ^[12]^. Evaluation of cells expressing EML4-ALK V3 indicated remarkably altered morphology and actin architecture, akin to that of the aNEK9- and aNEK7-expressing cells previously shown (Fig. S5A).

Upon depletion of cortactin with either of the two siRNAs, the elongated cytoplasmic protrusions were lost (Fig. 4A-C). The reduction of cytoplasmic protrusion length was more substantial upon depletion with siCTTNb than siCTTNa consistent with the more effective depletion with siCTTNb. Similarly to the effect in aNEK9 and aNEK7 cells, cortactin depletion led to a significant increase in linear F-actin stress fibres across the EML4-ALK V3 expressing cells (Fig. S6A, B).

**Figure 4.**
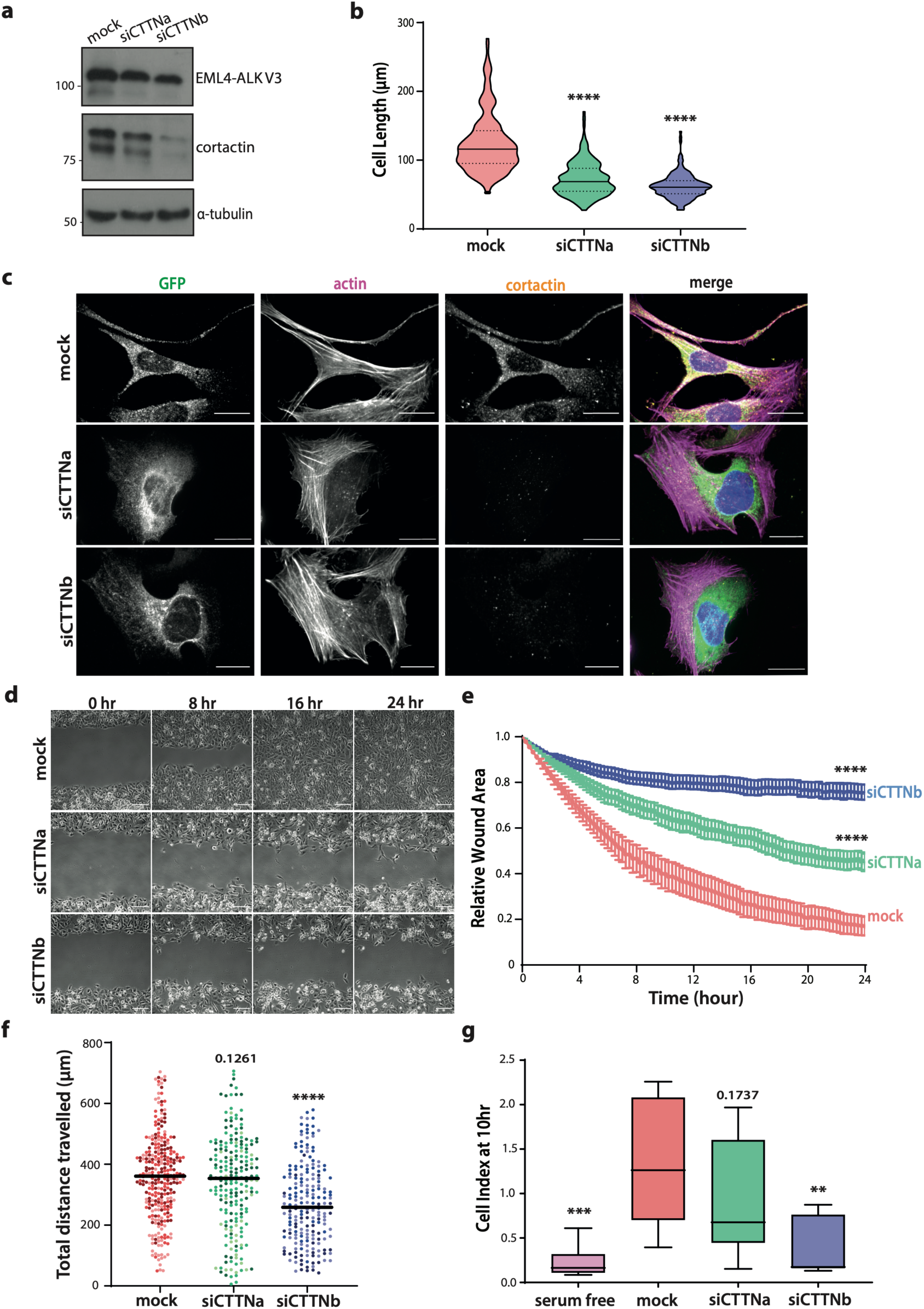
Cortactin is required for mesenchymal changes in cells expressing EML4-ALK V3. **A**. U2OS cells constitutively expressing YFP-EML4-ALK V3 were either mock depleted or depleted of cortactin using siCTTNa or siCTTNb for 48 hr. Cells were lysed and analysed by Western blot with ALK (EML4-ALK V3), cortactin and α-tubulin antibodies. M. wts (kDa) are indicated on the left. **B & C.** Cells treated as in A were stained with antibodies against GFP (YFP-EML4-ALK V3; green) and cortactin (yellow). Actin was stained with TRITC-phalloidin (magenta); merge images also show DNA stained with Hoechst 33258 (blue). Scale bars, 10 μm. Cell length measurements were determined as previously indicated and are shown in B. Solid lines indicate median and dotted lines the quartiles. **D.** For wound healing experiments, cells treated as in A were plated in a confluent single layer. A single scratch wound was made through the centre of each well and movement of cells measured by time-lapse microscopy over 24 hr. Representative images at time points indicated are shown. Scale bar, 100 μm. **E.** Based on data shown in D, the relative wound area over 24 hr with the mean and SEM are indicated. Final wound areas were analysed for significant differences (n=3 with 6 replicates per experiment). **F.** For single cell migration experiments, cells treated as in A were seeded on to collagen and single cell migration measured over 12 hr. Each shade of colour within plots represents a biological repeat. Mean and SEM are indicated (n=3 with 6 replicates per experiment). **G.** Cells treated as in A were subject to chemotactic migration experiments as described in Fig. 3D & H. The cell index at the 10 hr time point is indicated (n=3 with 2 replicates per experiment). **, *p<*0.01, ***, *p*<0.001, ****, *p*<0.0001 by one-way ANOVA.

To assess cell migration, we examined the consequences of cortactin depletion in the same three assays used previously that measure collective cell migration (wound healing assay), individual cell migration (single cell tracking) and directed cell migration (RTCA assay). In the wound-healing assay, depletion of cortactin caused a decrease in the rate of wound closure in cells expressing EML4-ALK V3, with the stronger depletion using siCTTNb causing almost complete abrogation of migration (Fig. 4D, E). Meanwhile, in single cell tracking assays, although siCTTNa only gave a weak and not significant reduction in migration rate, the more effective siCTTNb depletion led to significant impairment of migration with cells travelling a mean distance of 270 μm in 12 hr compared to 359 µm for mock-depleted cells (Fig. 4F). Finally, in the RTCA assay, cortactin depletion led to a reduction of directed cell migration, with siCTTNb again more effective that siCTTNa (Fig. 4G). Hence, this demonstrated that cortactin is required for changes in cell morphology and accelerated migration not only in cells expressing activated NEK9 and NEK7, but also cells expressing the EML4-ALK V3 fusion protein.

### Cortactin co-localises with EML4-ALK V3 and actin in filopodia-like extensions

To explore the localization of cortactin in cells expressing EML4-ALK V3, we switched to Beas-2B bronchial epithelial cells, a more relevant model system for NSCLC. Previous work had demonstrated that expression of EML4-ALK V3 in Beas-2B cells led to highly similar cellular phenotypes to those described earlier and that the cytoplasmic protrusions contained both actin and microtubules ^[12,41]^. Direct comparison between U2OS and Beas-2B phenotypes with and without expression of EML4-ALK V3 showed similar differences in morphology and both single and collective migration (Fig. S5A-D).

More detailed confocal microscopy analysis of Beas-2B cells expressing EML4-ALK V3 demonstrated that similarly to U2OS cells, upon depletion of cortactin the cytoplasmic protrusions were depleted (Fig. S7A-C). Interestingly, an exaggerated phenotype was observed upon depletion, whereby cells became larger with very distinct F-actin stress fibre networks detected across the cells. Corresponding with this, quantification of the actin filaments demonstrated increased length and decreased curvature with cortactin depletion (Fig.S7D-E).

Closer analysis of the cytoplasmic protrusions revealed that, as observed in cells expressing aNEK7, multiple fine FLEs were detected that contained not only actin but also EML4-ALK V3 and cortactin (Fig. 5A). Cortactin decorated the ends of FLEs and leading edges of protrusions. Quantification revealed that cortactin and actin co-localised within FLEs and at the branch-points to a significant extent, but not within the cell body (Fig. 5B). Meanwhile, EML4-ALK V3 colocalised with both actin and cortactin at the branch-points but not along the length of the FLEs or the cell body (Fig. 5B). These data suggest that EML4-ALK V3 might play an active role in the actin-branching activity of cortactin and actin-based remodelling of the plasma membrane.

**Figure 5.**
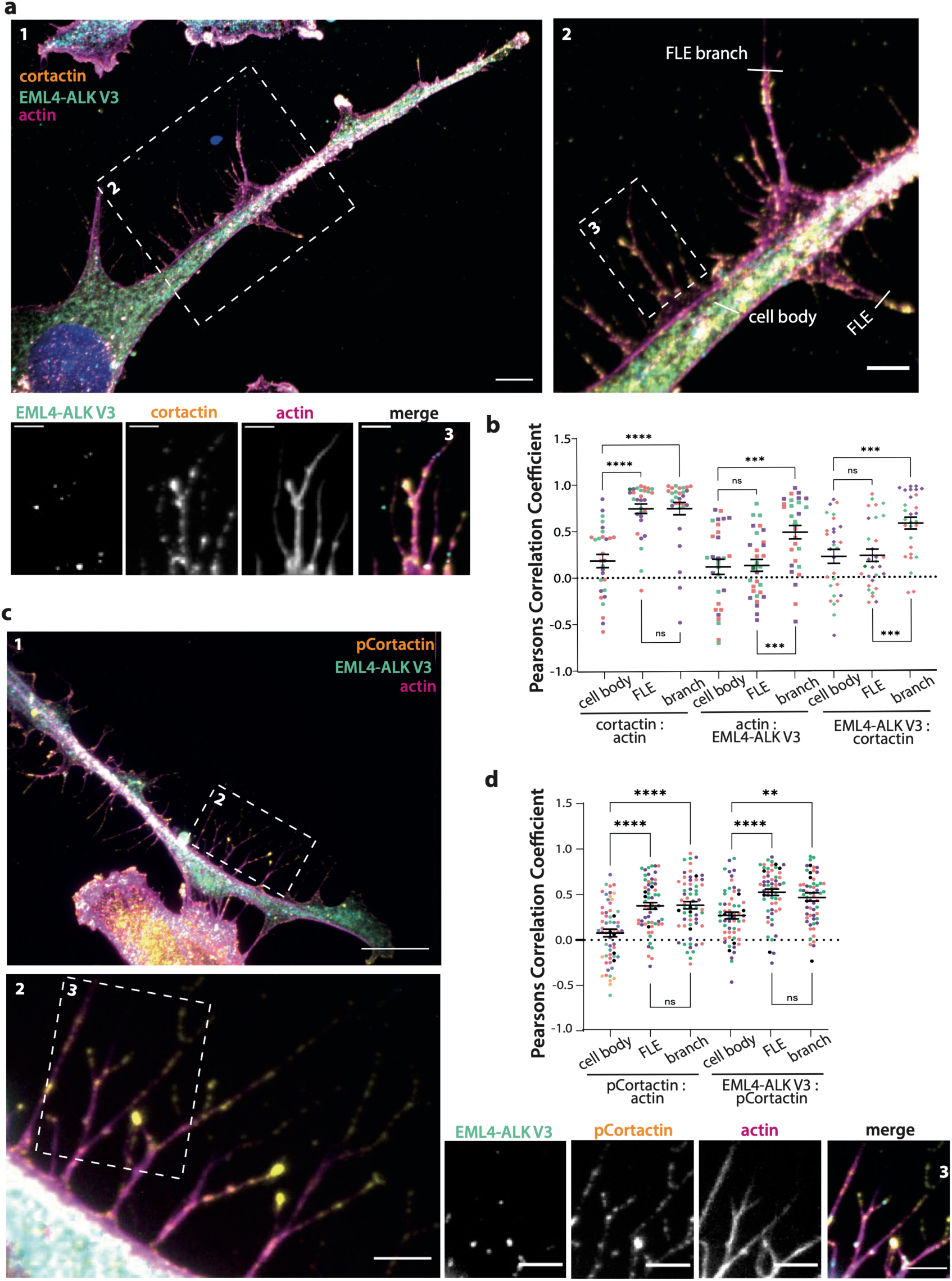
EML4-ALK V3, actin and phosphorylated cortactin colocalise in filopodia-like extensions. **A.** Beas-2B cells were induced to express YFP-EML4-ALK V3 for 72 hr before being fixed and stained with antibodies against cortactin (yellow) and YFP (EML4-ALK V3; cyan). Actin was stained with TRITC-phalloidin (magenta), while DNA was stained with Hoechst 33258 (blue). Magnified views are shown in panel (2) and (3), with the cell body, filopodia-like extensions (FLEs) and branch-points on the FLEs indicated. Scale bars, 10 μm (1) and 5 μm (2,3). **B.** Based on images as represented in A, 2.5 μm lines were drawn using Fiji software across actin fibres in the cell body, and across FLEs and FLE branches. The pixel intensities of cortactin, EML4-ALK V3 and actin were measured across the lines and Pearson’s correlation coefficient of colocalization calculated. Each different colour represents a repeat experiment (n=3). **C.** Beas-2B cells were induced to express EML4-ALK V3 before being fixed and stained with antibodies against phosphorylated cortactin (pCortactin, yellow) and YFP (EML-ALK V3, cyan). Actin was stained with TRITC-phalloidin (magenta), while DNA was stained with Hoechst 33258 (blue). Scale bars, 10 μm (1) 5 μm (2,3). **D.** Pearson’s correlation coefficient of colocalization was calculated for the protein combinations in the regions indicated as in C. Each different colour represents a repeat experiment (n=3). **, *p<*0.01, ****, *p*<0.0001 by one-way ANOVA.

Beas-2B cells expressing EML4-ALK V3 were probed with the pCortactin antibody previous used in initial *in vitro* phosphorylation assays. Strikingly, this revealed pCortactin colocalization with both actin and EML4-ALK V3 at the tips and branches of the FLEs (Fig. 5C & D). Together with the demonstration of colocalisation of phosphorylated cortactin and NEK7 at FLEs, this provides persuasive evidence that EML4-ALK V3, NEK7 and the associated phosphorylated cortactin subtype are enacting a role in formation of FLE, particularly at branch points.

### A phospho-mimetic cortactin mutant phenocopies expression of EML4-ALK V3 and activated NEK7

To examine how phosphorylation of cortactin in the ABR might influence the phenotypes described here, the conserved serine residues identified as phosphorylation sites in the first four F-actin binding repeats (Ser98, Ser135, Ser172 and Ser209) were mutated to either aspartic acid (S*4D) or alanine (S*4A) to create phospho-mimetic and phospho-null mutants, respectively, that could be expressed in mammalian cells. Additional silent mutations were introduced to make them RNAi-resistant to allow depletion of host cortactin and expression of mutant protein alone.

Together with a comparable WT cortactin construct, these were stably expressed with a C-terminal YFP tag in U2OS cells and cell morphology and actin organization examined by immunofluorescence microscopy (Fig. 6A, B). Cells expressing WT cortactin were similar in appearance to the parental cells albeit less cobblestone-shaped and exhibiting the presence of actin-containing filopodia. Cells expressing the phospho-mimetic S*4D cortactin protein exhibited more exaggerated versions of these changes, including more FLEs on the cell surface reminiscent of those observed in cells expressing activated NEK7 or EML4-ALK V3. It is noteworthy from the Western blot that there is increased expression of the phospho-mutant compared to wild-type proteins, hence the more pronounced phenotype could be due to this enhanced expression (Fig. 6A).

**Figure 6.**
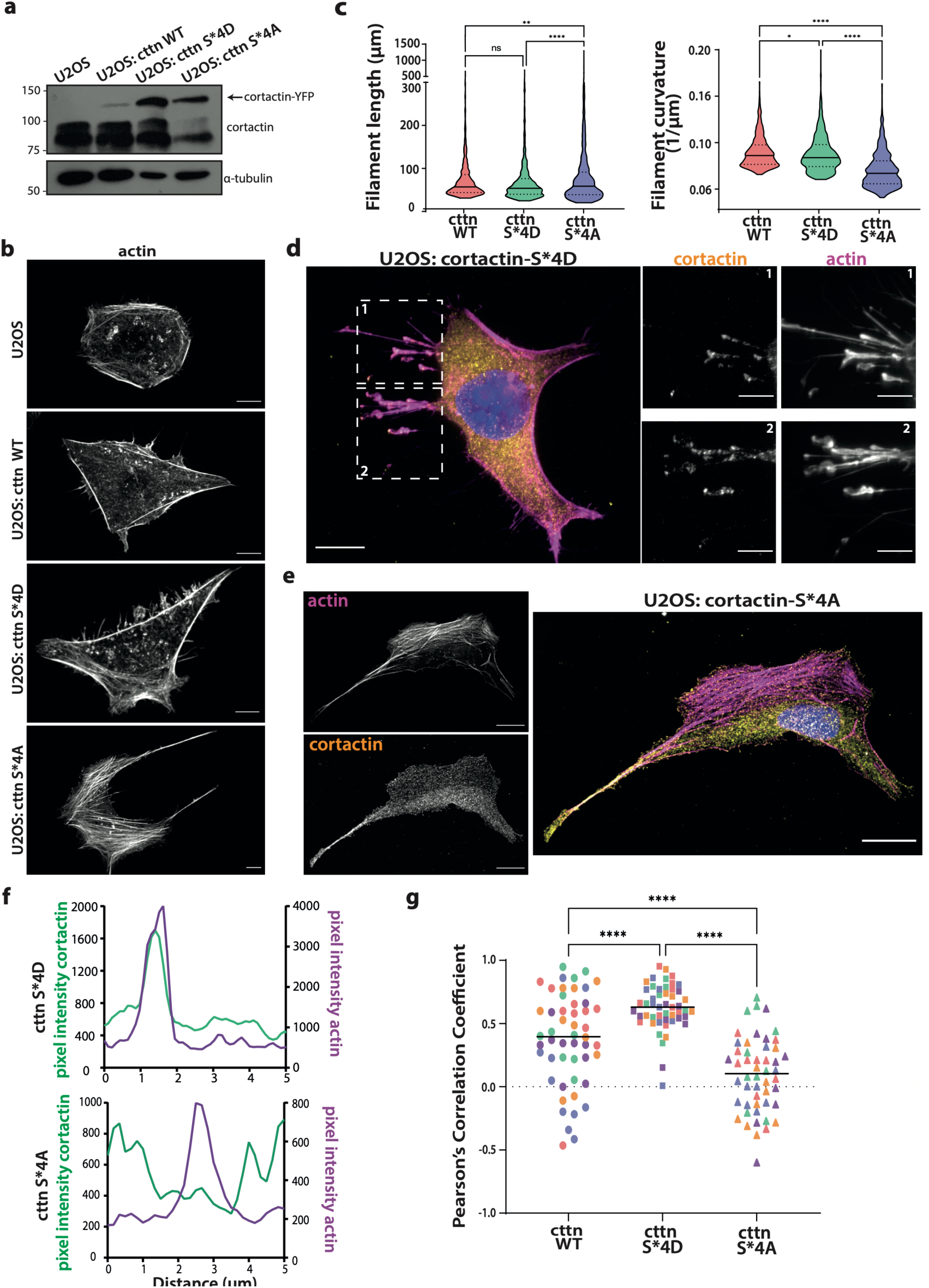
Expression of cortactin phospho-mutants alters cell morphology and actin organization. **A**. Parental U2OS cells or U2OS cells expressing YFP-tagged wild-type (WT), phospho-mimetic (S*4D) or phospho-null (S*4A) cortactin (cttn) were analysed by Western blot using cortactin and α-tubulin antibodies. M. wts (kDa) are indicated on the left. **B.** Cells as in A were stained with TRITC-phalloidin to visualise the actin cytoskeleton. Scale bars, 10 μm. **C.** Actin filaments of U2OS cells expressing cortactin-WT, -S*4D or -S*4A were analysed for length (left) and curvature (right). Violin plots include a solid line representing the median and dotted lines the quartiles (n=5 with 200 measurements per cell). **D.** Representative image of U2OS:cortactin-S*4D cell stained for actin (magenta), cortactin (yellow) and DNA (blue). Scale bar, 10 μm. Individual cortactin and actin stains showing the filopodia-like extensions from the two regions indicated are shown on right. Scale bars, 5 μm. **E.** Representative image of U2OS:cortactin-S*4A cell stained as in D. Scale bars, 10 μm. **F.** Representative colocalisation plots of cortactin and actin pixel intensity across a line of interest in U2OS cells expressing cortactin-S*4D or -S*4A. **G.** Pearson’s correlation coefficient of colocalisation indices of cortactin and actin in U2OS cells expressing cortactin-WT, - S*4D or -S*4A. Line indicates the mean and different colours individual experimental repeats (n=3 with 3 cells analysed per experiment and 10 individual 2.5 μm lines drawn over actin filaments). *, *p<*0.05, **, *p<*0.01, ****, *p*<0.0001 by unpaired Student’s T-test.

Strikingly, cells expressing the phospho-null (S*4A) mutant had a very different morphology. They frequently adopted a crescent-shape, with a wide convex leading edge and thin trailing edges (Fig. 6B). Live cell imaging confirmed that migration occurred in the direction of the leading edge (Movie S3). Actin organization was also different in cells expressing S*4A with an increased concentration of F-actin stress fibres that had a marginally increased length and substantially reduced curvature than cells expressing WT cortactin or the S*4D mutant (Fig. 6C). Increased actin stress fibres had also been observed simply upon cortactin depletion but without cells adopting this convex shape. Hence, while both cortactin depletion and expression of the S*4A mutant affected F-actin organization, the consequences on actin organization and cell morphology were not equivalent. No FLEs were seen on the surface of cells expressing the phospho-null S*4A mutant. This adds weight to the hypothesis that formation of these cellular structures results from activation of NEK7, downstream of EML4-ALK V3, and phosphorylation of cortactin on its actin-binding repeats.

The localization of the phospho-mimetic and phospho-null cortactin protein in these cells was analysed. This revealed that the phospho-mimetic S*4D mutant predominantly localized at the tips of the FLEs present in these cells where it colocalised with actin (Fig. 6D). In contrast, the S*4A protein localised strongly to what appeared to the trailing edges of cells in a manner that was largely mutually exclusive with localization of F-actin (Fig. 6E). Quantification confirmed the colocalization of the S*4D but not S*4A mutant protein with actin in these cells (Fig. 6F, G). This suggests that phospho-mimetic mutations of the conserved serine residues in the ABR increase affinity of cortactin for specific populations of F-actin, at least in some regions of the cell, whereas phospho-null mutations interfere with binding of cortactin with F-actin.

Finally, we assessed the consequences of expression of these phospho-mutant cortactin proteins on U2OS cell migration. In a scratch-wound assay, cells depleted of endogenous cortactin but expressing wild-type or the phospho-mimetic S*4D cortactin closed the wound marginally but significantly faster than mock-depleted parental U2OS cells. Intriguingly, cells expressing the phospho-null S*4A protein closed the wound substantially faster than cells subjected to other experimental treatments (Figure 7A-B, and Movie S4). Furthermore, in single cell tracking experiments, there was no significant difference in migration of cells expressing wild-type or S*4D cortactin as compared to control cells, whereas cells expressing the phospho-null S*4A mutant exhibited dramatically increased migration (Figure 7C). It was observed that cells migrated randomly with large displacement values, seemingly without contact inhibition and little retraction of leading or trailing edges (Figure 7D, and Movie S4).

**Figure 7.**
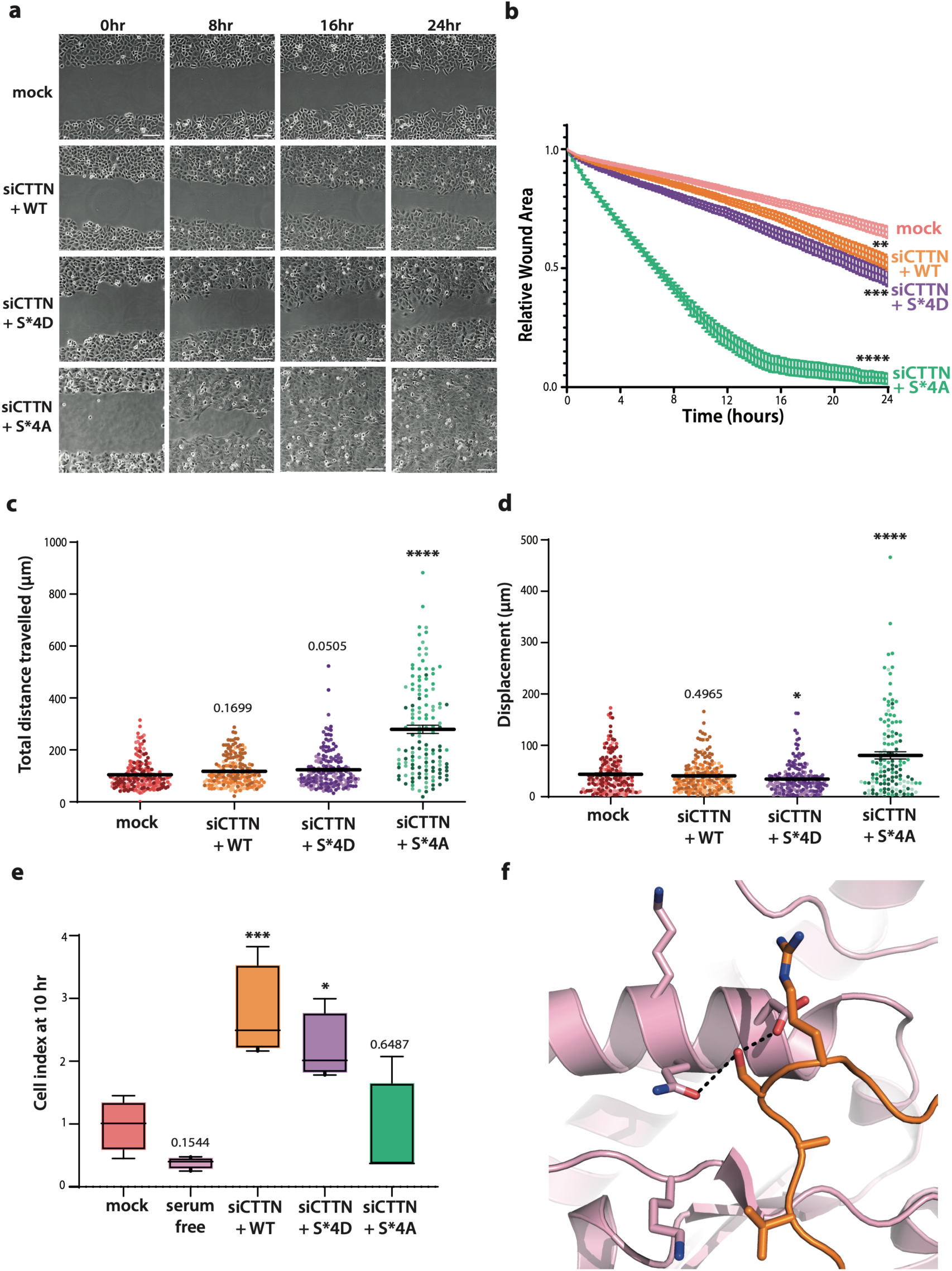
Expression of cortactin phospho-mutants alters cell migration. **A**. U2OS cells expressing recombinant cortactin-WT, -S*4D or -S*4A were depleted of endogenous cortactin using siCTTNb oligonucleotides for 72 hr. Parental U2OS cells (mock) and depleted cells were seeded in a confluent layer before a single scratch wound was made through the centre of each well and the movement of cells measured over 24 hr. Representative images at the time points indicated are shown from the wound healing assay. Cells were also fixed at the 8 hr time-point and stained with phalloidin-TRITC to visualise the actin cytoskeleton of the invasive front (bottom row). **B.** The relative wound area from cells treated as in A is indicated. Final wound areas were analysed for significant differences (n=3, 6 technical replicates per experiment). **C-D.** In single cell migration assays, cells treated as in A were plated onto collagen-coated wells and migration measured over 12 hr. Scatter plots showing the total distance travelled (C) and displacement (D) of cells are shown. Each shade of colour within plots represents a biological repeat. Mean and SEM are indicated (n=3 with technical 6 replicates per experiment). **E.** Chemotactic migration experiments were performed as described in Fig. 3D & H using cells treated as in A. The cell index at the 10-hr time point is indicated. Migration of mock-depleted cells with serum-free media was also measured. The mean and SEM are indicated (n=3 with 2 technical repeats per experiment). *, *p*< 0.05, **, *p*<0.01, ***, *p*<0.001, ****, *p*<0.0001 by one-way ANOVA. **F.** Molecular interaction of the first ABR of cortactin (orange) with an actin subunit (pink), in the vicinity of the Ser98 phosphorylated by NEK7. Two potential polar interactions are indicated with dashed lines. The image is based on the cryo-EM structure of Arp2/3 actin branch/cortactin complex (PDB entry 8P94)^27^.

Conversely, in the RTCA directional migration assay, cells expressing the WT or phospho-mimetic S*4D cortactin exhibited significantly increased migration compared to the control cells, whereas the phospho-null S*4A displayed minimal migration (Fig. 7E). This suggested that, despite the S*4A mutant displaying significant random migration, cells were not able to detect and migrate towards a chemo-attractant (directed migration). Therefore, we propose that phosphorylation on the ABR increases directional migration, perhaps through the action of FLEs, whereas the inability to phosphorylate this region leads to an accelerated but less directional form of cell migration. We therefore conclude that phosphorylation of cortactin within its ABR is likely to be an important mechanism through which cells expressing EML4-ALK V3 or activated NEK7 control changes to both cell morphology and cell migration.

## DISCUSSION

The successful control of the cytoskeleton is multifaceted, with numerous different signalling factors, proteins and post-translational modifications participating in highly complex and dynamic interactions ^[42,43]^. These events allow constant remodelling of the cytoskeleton that enable the cell to efficiently navigate between different environments and fulfil specific functions in both normal physiological and pathological settings. In cancer, cells gain the ability to migrate away from their primary site of origin and disseminate to distant sites in the process of metastasis. We have previously shown that the EML4-ALK V3 fusion protein, a major oncogenic driver in NSCLC, localises to and stabilises microtubules and induces a shift to mesenchymal-like properties that include altered cell morphology and increased directional migration ^[12]^. Moreover, we found that these phenotypes depend upon microtubule recruitment and activation of the NEK9 and NEK7 kinases. Here, we demonstrate that this signalling pathway is also dependent on the actin-binding and stabilising protein, cortactin, with our data pointing towards cortactin being a direct downstream substrate of NEK7. Hence, this study highlights a novel mechanism of interaction between cytoskeletal proteins and kinases that is potentially important for the increased metastatic potential of tumours that express the EML4-ALK V3 fusion.

Cortactin is implicated in multiple roles across both healthy and diseased cells in the lung and beyond, including in cell motility in aggressive cancers ^[23,25,44]^. Cortactin interacts with several other microtubule-associated proteins to facilitate actin-microtubule crosstalk in directed migration and invasion, including microtubule plus-end binding (EB) proteins, EB1 and EB3, as well as TIP150 and MAP1B^45–50^. Together, there is mounting evidence that cortactin is protein linchpin which both promotes the stabilisation and branching of actin via its N-terminal end, whilst interacting with other cytoskeleton-related proteins via its C-terminal end to coordinate cytoskeletal and cellular dynamics.

The majority of previously identified phosphorylation events related to cortactin function are located in the proline-rich region, including those catalysed by SRC, ERK1/2, PAK, ABL and ARG ^[31,51–53]^. In this study identified NEK7 sites were spread across the NTA, ABR, α-helical and proline-rich regions. Shared NEK6/7 phosphorylation sites were found in sites within the ABR and proline-rich regions, with the previously characterised ERK1/2 phosphorylation site Ser405 of note ^[31,36]^. Further work is required to understand these phosphorylation events, particularly whether the shared sites between the two NEK kinases have similar functions. For example, as both NEK6 and NEK7 display enhanced activity during mitosis and cortactin-mediated actin branching is required in certain stages of mitosis, it is possible that the phosphorylation of cortactin by these kinases may also play a role in actin-mediated events during normal cell division ^[16,18,54]^.

In this study, we chose to focus on four phosphorylation sites -Ser98, Ser135, Ser172 and Ser209 – that reside in a conserved sequence within the first four ABR repeats. These repeats are important for cortactin function with Ser209 essential for actin binding and stabilisation ^[27,35,55–57]^. The recent cryo-EM structure of cortactin bound to an actin branch reveals the critical role of the ABR repeats in binding actin at the branching filament (Fig. 7F) ^[27]^. The first ABR repeat was resolved and the conserved Ser side chain makes polar interactions with actin, which would be disrupted by mutation to Ala. This aligns with data shown here where cells expressing cortactin S*4A showed significantly altered phenotypes both in actin structure and resulting migration. The effect of phosphorylation of the Ser on residue interaction is less clear, but there are several basic residues nearby that could potentially form interactions with this modified residue. Intriguingly, acetylation of lysine residues 11-amino acids downstream of these four sites has been reported to reduce actin binding ^[40]^. It is therefore tempting to speculate that different combinations of phosphorylation and acetylation in the ABR could result in cortactin proteins with a distinct range of actin binding affinities, perhaps fine-tuned to different spatiotemporal contexts.

Our data suggests that cortactin is a downstream target of NEK7 in cells expressing EML4-ALK V3 to facilitate the most effective spatiotemporal alteration of actin filaments alongside that of microtubules. The coordination of the cytoskeleton is crucial to alter the properties of cells in such a way that increases the rate of effective migration ^[42,58,59]^. Long cytoplasmic protrusions and accelerated migration are characteristics of EML4-ALK V3 expressing cells. Almost identical mesenchymal-like phenotypes are observed in cells expressing constitutively active NEK9 and NEK7. Upon depletion of endogenous cortactin, these phenotypes were reversed, whereby cells became cobblestone-shaped with reduced migration. This demonstrates the requirement for cortactin for the mesenchymal-like changes driven by the EML4-ALK V3:NEK9:NEK7 pathway.

It could be argued that the actin branching activity of cortactin is simply required to provide the robust cellular architecture that is necessary for these phenotypes and that this alone is not evidence that cortactin is a downstream target of this signalling pathway. However, the fact that similar phenotypes were observed in cells expressing a cortactin protein with phospho-mimetic mutations in sites shown to be phosphorylated by NEK7, together with the spatially restricted co-localization of these proteins in migratory structures in these cells, strongly suggests a more direct link.

In our study, it was observed that striking actin stress fibre structures were formed both upon expression of phospho-null cortactin and depletion of cortactin in the EML4-ALK, activated NEK9 and NEK7 cell models. Previous studies have demonstrated the role of cortactin through depletion and subsequent abrogation of functional invadopodia, podosomes or dendrite formation ^[60–62]^. However, there is evidence to suggest a link between cortactin deficiency in endothelial cells, the formation of stress fibres through activation of the ROCK1-mediated actomyosin contractility pathway and endothelial barrier dysfunction ^[26,63]^. As Rho-ROCK activity supports amoeboid migration in cancer cells, it is intriguing to hypothesise whether the enhanced random migratory phenotypes and lack of directional migration observed in phospho-null cortactin expressing cells is due to a cortactin-induced switch to ROCK signalling, similar to that observed in endothelial cells, however further work is required to support this hypothesis.

In another intriguing observation, cells expressing EML4-ALK V3, active NEK7 or phospho-mimetic cortactin all produced FLEs along the surface of cytoplasmic protrusions. Interestingly, these are reminiscent of dendritic spines in neuronal cells. The distribution and function of both cortactin and NEK7 have been separately implicated in modulating dendritic spine morphology in neurons through alteration of microtubule and actin dynamics ^[47,48]^. The FLEs identified in our study differed from classical filopodia as they were often branched and contained “bulbs” at the ends of the extensions, which were shown to contain actin, cortactin, EML4-ALK V3 and activated NEK7. Importantly, phosphorylated cortactin was also found colocalised at the ends and branches of the FLEs, further highlighting the role of the EML4-ALK:NEK9:NEK7 module in spatially controlling the activity of cortactin to build these structures. Although the role of FLEs in these cells is unclear, similar structures - described as ‘migrasomes’ - have been implicated in cell-cell communication and cell migration through their release from migrating cells ^[64,65]^. Consistent with our hypothesis that phosphorylated cortactin is required for these structures, cells expressing phospho-null cortactin did not form FLEs and did not migrate in a chemotactic manner. Chemotaxis is a crucial step of tumour cell dissemination. Therefore, it is likely that in highly metastatic EML4-ALK V3 NSCLC tumour cells this pathway enables not only an increased rate but also highly directed form of migration.

In summary, these findings demonstrate the requirement of cortactin to enable directed and effective migration of cells expressing EML4-ALK V3 and identifies cortactin as a novel target of NEK7. For cells to migrate effectively, highly dynamic protrusive cytoplasmic structures are required, which in turn require fine-tuning and control of cytoskeletal filaments and associated proteins. EML4 is highly expressed in neuronal cells, while ALK is predominately expressed during development of the nervous system ^[66,67]^. Meanwhile, as indicated above, cortactin and NEK7 have both been implicated in formation of neuronal dendritic spines that resemble the FLEs described in this study ^[47,48,62]^. This raises the tantalizing prospect that ALK, EML4, NEK9, NEK7 and cortactin could cooperate in a normal physiological mechanism to control cytoskeletal dynamics in neuronal cells but that this process has been hijacked by EML4-ALK V3 expressing cells to promote cancer cell migration. Further research is required to map and characterise these interactions in more detail. Meanwhile, this work provides a step forward in understanding the complex web of protein interactions and signalling during migration.

## MATERIALS & METHODS

### KESTREL analysis and mass spectrometry

The KESTREL screen that led to identification of cortactin as a NEK6 substrate was performed as described in O’Regan *et al.* (2015) ^[18]^. Briefly, cell lysates of HEK293 cells were prepared, insoluble material removed, and supernatant filtered before being degassed and desalted. The desalted extract was applied to a 25 ml Heparin Sepharose High Performance column. During elution, forty 10 ml fractions were collected. 20 µl of each fraction was diluted 1:10 in KESTREL test buffer (50 mM Tris.HCl, pH 7.5, 7 mM β-mercaptoethanol, 1 mM EGTA, 10 µg/ml leupeptin, and 1 mM Pefabloc) and incubated for 5 min with 3 mM MnCl_2_ and 1 kBq/ml [^32^P]-γ-ATP in the absence or presence of 1 µg/ml active recombinant NEK6 (Invitrogen). Reaction products were analyzed by SDS-PAGE and subsequent autoradiography. Fractions identified to contain NEK6 substrates were pooled, desalted and further separated along a 10 ml gradient to 1 M NaCl on a 1 ml Source 15Q column, and analyzed. Substrate containing fractions 16–18 were pooled, separated by size using a 120 ml Superdex 200 column, and analyzed. Superdex 200 fractions 3–7 were pooled, concentrated by filtration, and incubated with either 3 mM MnCl_2_ or 10 mM Mg-acetate in the presence of [^32^P]-γ-ATP, with or without 1 µg/ml NEK6. The reactions were denatured, alkylated with 50 mM iodoacetamide, and separated by SDS-PAGE, stained and subsequently analysed. Protein bands that had been radiolabeled in the presence of NEK6 were excised from the gels, destained, digested with trypsin, and subjected to mass spectrometry fingerprinting (matrix-assisted laser desorption/ionization–tandem time of flight) using a proteomics analyzer (4700; Applied Biosystems) for data acquisition and the Mascot search engine (Matrix Science) to search for matches in the NCBInr database.

### Identification of proteins and phosphorylation sites by Mass Spectrometry

For protein and phosphorylation-site identification samples were separated via SDS-PAGE electrophoresis, stained with Coomassie blue and gel pieces subjected to an in-gel digestion: gel pieces were washed twice in 50% Methanol, 0.1 M triethylammonium bicarbonate (TAB) pH 8.0, then alkylated with 50mM iodoacetamide, 2mM TCEP, 0.1M TAB, then desiccated in 100% Methanol and then digested by incubating with 5 μg/ml trypsin in 50 mM TAB (Sigma-Aldrich) overnight. Supernatants were collected, dried in a Speed-Vac, resuspended in 30 μl 0.1 % formic acid and subjected to liquid chromatography–mass spectrometry (LC-MS) analysis using an Ultimate 3000 RSLC nano system coupled to Exploris 480 mass spectrometer (ThermoFisher Scientific) 5 µl samples were injected and peptides were loaded onto a nanoViper C18 Trap column (5 µm particle size, 100 µm x 2 cm) and separated in a C18 reversed phase Easy-spray column (2 µm particle size, 75 µm x 50 cm) (ThermoFisher Scientific) at a flow rate of 300nl/min. A linear gradient was used, starting at 3% B and maintained for 2 min, from 3-7% B in 5min, 7-25% B for 43 min, 25-35% B for 5 min, 35-95 B over 3min, maintained at 95% B for 2 min, 95-3% B in 1 min and maintained at 3% B for 9 min. Solvents used were A: 0.1% formic acid and B: 80% acetonitrile (ACN) with 0.08% formic acid.

Mass Spectrometry data was acquired in data-dependent mode using the following parameters: MS1 spectra were acquired in the Orbitrap at a resolution of 60,000 (at 400 m/z) for a mass range of 350-1200 m/z with a FTMS full AGC target of 300%. The most intense ions (with a minimal signal threshold of 10000) were selected for MS2 analysis and were fragmented (using CID with a collision energy of 35%), and then dynamically excluded for 30 seconds.

Data was analysed using Proteome Discoverer v.2.4 and Mascot search engine using Swiss Prot database with taxonomy restricted to human proteins (20,402 sequences). Parameters used were the following: Variable modifications: Oxidation (M), Dioxidation (M), Phospho (STY); Fixed modifications: Carbamidomethyl (C), Enzyme: Trypsin/P, Maximum missed cleavages: 3, Precursor tolerance: 10ppm, MS2 tolerance: 0.6Da, Minimum score peptides: 18. Phospho-site assignment probability was estimated via Mascot and ptmRS (Proteome Discoverer v.2.4).

### Plasmid construction and mutagenesis

YFP-tagged EML4-ALK variants, NEK7-Y97A (aNEK7) and NEK9-ΔRCC1 (aNEK9) constructs were generated as previously described ^[12]^. Full-length cortactin with a YFP tag was obtained from Addgene. RNAi resistant cortactin phospho-mutant constructs, S*4D and S*4A, were generated using QuickChange II XL Site-Directed Mutagenesis Kit (Agilent Technologies). Following digestion of methylated parental DNA, plasmids were transformed into competent *E. coli* DH5α cells. Colonies were selected and expanded before DNA was isolated using ZymoPURE™ II Plasmid Midiprep Kit (Zymo Research). Resulting DNA was sequenced by Source Bioscience (Nottingham) to confirm site-specific mutagenesis.

### Protein production

To complete kinase assay time course screens of NEK6 and NEK7 phosphorylation of cortactin, cortactin A was produced as a 6-His-Halo-TEV-Cortactin A fusion protein (Dundee: DU77714) in BL21 DE3 cells, purified over Ni-NTA agarose, cleaved of the tag and polished over a Superdex 200 column. Recombinant 6-His-tagged NEK6 or NEK7 were initially purchased from Invitrogen and then from MRC-PPU reagents (mrcppureagents.dundee.ac.uk), where NEK6 is DU635 and NEK7 is DU631.

To determine phosphorylation events occurring on WT or S*4D cortactin protein, cortactin WT or S*4D plasmids were used to transform Rosetta™(DE3) competent cells (Novagen) on chloramphenicol/ampicillin agar plates. Single colonies were expanded in chl/amp TY buffer at 37°C for 5-6 hr until the optical density was 0.1. Protein production was induced using 40 μM IPTG at 20°C for 16 hr. The cultures were collected, pelleted at 4000 rpm at 4°C for 10 mins, lysed and sonicated. The lysed cells were centrifuged at 30000 rpm at RT for 20 mins and the supernatant collected. His-Pur Ni-NTA beads (Fisher) were washed and incubated with the supernatant at 4°C for 30 mins. The mix was subsequently centrifuged and washed three times at 4000 rpm at 4°C for 2 mins before being resuspended in elution buffer (50 mM Tris/HCl pH8.0, 500 mM NaCl, 300 mM Imidazole pH8.0, 10% glycerol, 1 mM DTT) at 4°C for 15 mins. After centrifugation at 4000 rpm at for 2 mins the supernatant was collected and stored at -80°C.

### Phosphoantibody production

Phospho-specific cortactin (pCortactin) antibodies were generated in rabbits against residues 166-179 with a modified phosphorylated S172 (Peptide Specialty Laboratories, GmbH). Antibodies were purified by sequential passage over a column containing the non-phosphorylated peptide and subsequently a column containing the phosphorylated peptide. Columns were washed extensively with 10 mM sodium phosphate, pH 6.8, and specific antibodies were eluted with 0.1 M glycine, pH 2.4, containing neutralizing quantities of 2 M K_2_HPO_4_.

### Kinase assays

To complete kinase assay time course screens of NEK6 and NEK7 phosphorylation of cortactin, 4µg of protein, equivalent of 3 µg of recombinant human cortactin A, was incubated with 0.1 µg of human His-NEK6 or human His-NEK7 in the presence of 10 mM MgCl_2_ and 0.1 mM [γ^32^P]ATP. The reactions were terminated by adding SDS-loading buffer and incubation at 90°C for 5 min. The samples were separated on 4%-20% Tris-Glycine SDS-gels and stained with Coomassie BlueInstant Blue. After destaining, the gels were scanned and then used to expose X-Ray films (Cytiva). The gel pieces representing each time point were excised and decay was measured in a Scintillation Counter. For calibration and calculations 1 µl aliquots, equivalent to 1 nmol of the [γ^32^P]ATP solutions were used.

To determine phosphorylation events occurring on WT or S*4D cortactin protein, 0.1 μg of purified NEK6 or NEK7 (Millipore) were pre-incubated in kinase buffer (50 mM HEPES.KOH pH7.4, 5 mM MnCl_2_, 5 mM β-glycerophosphate, 5 mM NaF, 10 mM ATP, 1 mM DTT) for 1 hr prior to assay start. The kinases were mixed with 25 μg of cortactin WT or S*4D purified protein and 90 mM ATP and incubated at 30°C for 6 hr. The reactions were terminated by adding 3X Laemmli sample buffer and the samples heated to 90^0^C for 5 min. Samples were separated on an SDS-PAGE gel and analysed by Western blot.

### Cell culture, drug treatments and transfection

HeLa, U2OS and derived stable cell lines were grown in Dulbecco’s Modified Eagle’s Medium (DMEM). Beas-2B cells were grown in Roswell Park Memorial Institute (RPMI) 1640 Medium. All media was supplemented with 10% heat-inactivated fetal bovine serum (FBS), 100 U/ml penicillin, and 100 µg/ml streptomycin, at 37°C in a 5% CO_2_ atmosphere. U2OS cells constitutively expressing YFP-EML4-ALK V3 were generated as previously described^12^ and maintained using media supplemented with 100 µg/ml hygromycin (Invitrogen). Doxycycline-inducible U2OS:NEK9-ΔRCC1 and HeLa:YFP-NEK7-Y97A cell lines were generated as previously described^12^ and maintained using media supplemented with 1 µg/ml of G418 and 800 ng/ml of puromycin (NEK9) or 200 µm/ml hygromycin B (NEK7). Beas--2B cells were induced to express EML4-ALK V3 upon doxycycline treatment as described previously ^[12]^. Doxycycline was used to induce expression of constructs at a final concentration of 1 µg/ml for 72 hr (unless otherwise stated). All cell lines were cultured for no more than 20 generations and were routinely tested for mycoplasma contamination using an in-house PCR test.

To generate stable cortactin WT-, S*4D- and S*4A-expressing cell lines, U2OS cells at 40% confluency were transfected with the appropriate plasmids using FuGENE transfection reagent (Promega) at a 3:2 ratio (FuGENE:DNA). After 48 hr, cells were collected and analysed by flow cytometry using the BD FACSCanto™II (BD). Single cells showing expression of FITC were sorted into a 96-well plate with high FBS containing RPMI media with one cell per well. The single cell clones were grown in selection media and tested for expression by Western Blot and immunofluorescence microscopy.

For transfection and RNAi depletion experiments, cells were seeded at 40% confluency in OptiMEM media. For transfection, FuGENE transfection reagent was mixed with DNA plasmids at a 3:2 ratio. For RNAi depletions, 100 nmol/L siRNA oligonucleotides and oligofectamine (Invitrogen) were combined in OptiMEM according to manufacturer’s instructions. Seeded cells were washed twice with FBS-free OptiMEM, and the mixtures added to cells in a dropwise manner and incubated at 37°C 5% CO_2_ for 4 hr. After 6 hr, FBS was added to the cultures so the final concentration was 10% (v/v) and incubated a further 48-72 hr before analysis. siRNA oligonucletoides directed against cortactin were s4665 (siCTTNa) and s4666 (siCTTNb) (Thermo Fisher Scientific, #4392420, #4392420).

### Preparation of cell extracts, SDS-PAGE and Western blotting

Cells were lysed in RIPA lysis buffer (50 mM Tris-HCl pH 8, 150 mM NaCl, 1% v/v Nonidet P-40, 0.1% w/v SDS, 0.5% w/v sodium deoxycholate, 5 mM NaF, 5 mM β- glycerophosphate, 30 µg/ml RNase, 30 µg/ml DNase I, 1x Protease Inhibitor Cocktail, 1 mM PMSF) prior to analysis by SDS-PAGE and Western blotting. Primary antibodies were: cortactin 4F11 (1:2000, Millipore, #05-180), GFP (1:2000, Abcam, #ab290), α-tubulin (1:5000, Sigma, #T5168), myc (1:1000, Cell Signalling Technologies, #2276), NEK9 (1:200, Santa Cruz Biotechnology, #sc-100401), NEK7 (1:200, Aviva Systems, #OAEB01009), EML4 (1:500, Thermo Fisher, #A301-908A), ALK (1:1000, Cell Signalling Technology #3633), and GAPDH (1:2000, Cell Signalling Technology, #2118). Secondary antibodies used were anti-rabbit, anti-mouse or anti-goat horseradish peroxidase (HRP)-labelled IgGs (1:2000; Sigma, AP160P, 12-349, AP106P). Western blots were detected using enhanced chemiluminescence (Pierce).

### Fixed and time-lapse microscopy

Cells grown on acid-etched glass coverslips were fixed with 3.7% formaldehyde at 37°C for 20 min. After permeabilization with 0.5% Triton X-100 for 10 mins, cells were blocked in PBS with 3% BSA. Primary antibodies used were against cortactin (1:2000, Millipore, #05-180), GFP (0.5 µg/ml, rabbit; Abcam, #ab290), α-tubulin (0.1 µg/ml, mouse, Sigma, #T5168), myc (1:1000, mouse, Cell Signalling Technologies, #2276), ALK (1:1000, Cell Signalling Technology #3633) and NEK7 (0.8 µg/ml, goat, Aviva Systems, #OAEB01009). Secondary antibodies were Alexa Fluor 488, 594 and 647 donkey anti-rabbit, donkey anti-mouse and donkey anti-goat IgGs (1 µg/ml; Invitrogen, #A-21206, 21206, 11055 21207, 21203, 11058, 31571, 31573, 21447). DNA was stained with Hoechst 33258 (Thermo Fisher, #H3569) and actin with TRITC-phalloidin (BioTechne, #5783). Imaging was performed on a VisiTech infinity 3 confocal microscope fitted with Hamamatsu C11440 -22CU Flash 4.0 V2 sCMOS camera and aPlan Apo VC 60x (NA 1.4) or Plan Apo 100 x objective (NA 1.47). For super-resolution radial fluctuations (SRRF) microscopy, 100 images from the same slice were captured per channel and processed using the nanoJ-SRRF plugin in Fiji ^[68]^.

To quantify cell morphology, images of cells stained with TRITC-phalloidin (actin) and Hoescht 33258 (DNA) were analysed using Fiji^69^. Actin and DNA images were merged and the following parameters were measured: (i) cell length: a line was drawn at ∼180° between the furthest two points of the cell and which passed through the nucleus and the distance measured; (ii) cell width: a line was drawn at ∼180° between the closest two points of the cell which passed through the nucleus and the distance measured; (iii) cell area: a free-hand shape was drawn around the cell perimeter and the area measured. To determine F-actin microarchitecture, images of cells stained with TRITC-phalloidin (actin) were analysed using SOAX ^[70]^. The first 250 highest values for snake length and curvature were analysed using GraphPad Prism.

To quantify co-localization on actin filaments, using Fiji a 2.5 μm line was drawn along actin filaments in the TRITC-channel of unmerged images. The intensities of actin, NEK7, EML4-ALK, cortactin and pCortactin along the line were quantified in Image/JFiji using ImageJ macro (Macro_plot_lineprofile_multicolor from Kees Straatman, University of Leicester, Leicester, UK) and the Pearson’s correlation coefficient analysed in Microsoft Excel.

Differential interference contrast and time-lapse microscopy were carried out using a Nikon eclipse Ti inverted microscope using a Plan Fluor 10x DIC objective (NA 0.3). Images were captured using an Andor iXonEM+ EMCCD DU 885 camera and NIS Elements software (Nikon). For time-lapse imaging, cells were cultured in 6-well dishes and maintained on the stage at 37°C in an atmosphere supplemented with 5% CO_2_ using a microscope temperature control system (Life Imaging Services, Basel, Switzerland) with images acquired every 15 min for 24 hr (migration assays) or every 10 sec for a 5 min period (filopodia observation).

### Cell migration assays

For individual cell tracking assays, cells were seeded at 30% confluency in 1 μg/ml collagen-coated 6-well plates. After 12 hr, cells were incubated in the Livecyte 2 imaging system (Phasefocus, Sheffield, UK) and 6 areas per well were imaged every 15 min for 24 hr using quantitative phase imaging and PLN 10x/NA=0.25 objective. Migration paths over 12 hr were analysed using TrackMate in ImageJ/Fiji ^[71]^.

For scratch-wound assays, cells were grown to 95% confluency in 6 well dishes before a pipette tip was used to scrape a 0.5 - 1 µm line across the width of the well. Cells were washed 3-5x in pre-warmed media and imaged by time-lapse microscopy. Movement of cells into the wound was observed through capturing images every 15 min in 6 areas per well over a 24 hr period using a Nikon Eclipse Ti microscope installed with Andor iXonEM+ EMCCD DU 885 camera and Plan Fluor 10x DIC objective (NA=0.3). Movement was analysed using the *MRI_Wound_Healing_Tool* macro from Montpellier Resources Imagerie on ImageJ/Fiji (https://github.com/MontpellierRessourcesImagerie/imagej_macros_and_scripts/wiki/).

Directed cell migration in real time was analysed using the xCELLigence Real-Time Cell Analyzer (RTCA) DC equipment (ACEA Biosciences) and CIM-16 plates, a 16 well system where each well is composed of upper and lower chambers separated by an 8 μm microporous membrane. Cells were grown in serum free medium for 24 h before being seeded in serum free medium into the upper chamber; complete medium containing 10% FBS was used as a chemo-attractant in the lower chamber of the plate and migration determined as the relative impedance change (cell index) across microelectronic sensors integrated into the membrane. Measurements were taken every 15 mins for 48 hr.

### Statistical analysis

All quantitative data represent means and SEM of at least three independent experiments, unless otherwise stated. Statistical analyses on data shown in histograms or dot plots were performed using a one-tailed unpaired Student’s *t* test assuming unequal variance or a one-way analysis of variance followed by post hoc testing for analysis of 2 data sets and multiple data sets, respectively. *p*-values represent: *, *p*<0.05; **, *p*<0.01; ***, *p*<0.001; ****, *p*<0.0001. n.s., non-significant.

## ACKNOWLEDGMENTS

We dedicate this publication to the memory of Prof Andrew Fry, who led the research and design of this study, as well as inspiring countless scientists to embrace adventure and pursue knowledge.

We thank the University of Leicester Core Biotechnology Services for providing access to the Advanced Imaging Facility (RRID:SCR_020967), Flow Cytometry and Protex Plasmid Cloning Service. We are grateful to Jene Choi (Asan Medical Centre, Seoul) for providing Beas-2B cells expressing EML4-ALK V3 and acknowledge support from undergraduate project students, Elinor Pulman and Lottie Grey-Wilson, for contributing to the mutagenesis of cortactin constructs.

## AUTHOR CONTRIBUTIONS

ELR and AMF conceived and planned the experiments. ELR and AK carried out the experiments with support from KRS, RG, DL, RET, TH and SWR. ELR wrote the manuscript with support from AMF and RB. AMF supervised the project with support from LO and RB. All authors helped shape the research and data analysis and provided critical feedback on the manuscript.

The authors declare that they have no conflict of interest.

## DATA AVAILABILITY STATEMENT

The datasets generated during and/or analysed during the current study are available from the corresponding author on reasonable request.

## REFERENCES

1. Sung, H. et al. Global Cancer Statistics 2020: GLOBOCAN Estimates of Incidence and Mortality Worldwide for 36 Cancers in 185 Countries. CA. Cancer J. Clin. 71, 209–249 (2021).

2. Tamura, T., et al. Specific organ metastases and survival in metastatic non-small-cell lung cancer. Mol. Clin. Oncol. 3, 217 (2014).

3. American Cancer Society. Lung Cancer Survival Rates | 5-Year Survival Rates for Lung Cancer | American Cancer Society. https://www.cancer.org/cancer/types/lung-cancer/detection-diagnosis-staging/survival-rates.html.

4. NHS England Digital. Cancer Survival in England, cancers diagnosed 2016 to 2020, followed up to 2021 - NHS England Digital. https://digital.nhs.uk/data-and-information/publications/statistical/cancer-survival-in-england/cancers-diagnosed-2016-to-2020-followed-up-to-2021 (2023).

5. Soda, M. et al. Identification of the transforming EML4–ALK fusion gene in non-small-cell lung cancer. Nature 448, 561–566 (2007).

6. Sabir, S. R., Yeoh, S., Jackson, G. & Bayliss, R. EML4-ALK variants: Biological and molecular properties, and the implications for patients. Cancers vol. 9 (2017).

7. Young, L. C. et al. Identification of novel isoforms of the EML4-ALK transforming gene in non-small cell lung cancer. Cancer Res. 68, 4971–4976 (2008).

8. Christopoulos, P., et al. *EML4-ALK* fusion variant V3 is a high-risk feature conferring accelerated metastatic spread, early treatment failure and worse overall survival in ALK ^+^ non-small cell lung cancer. Int. J. Cancer 142, 2589– 2598 (2018).

9. Elshatlawy, M., Sampson, J., Clarke, K. & Bayliss, R. EML4-ALK biology and drug resistance in non-small cell lung cancer: a new phase of discoveries. Mol. Oncol. 17, 950–963 (2023).

10. Qin, Z. et al. Phase separation of EML4-ALK in firing downstream signaling and promoting lung tumorigenesis. Cell Discov. 7, (2021).

11. Sampson, J., Richards, M. W., Choi, J., Fry, A. M. & Bayliss, R. Phase-separated foci of EML4-ALK facilitate signalling and depend upon an active kinase conformation. EMBO Rep. 22, (2021).

12. O’Regan, L. et al. EML4-ALK V3 oncogenic fusion proteins promote microtubule stabilization and accelerated migration through NEK9 and NEK7. J. Cell Sci. jcs.241505 (2020) doi:10.1242/jcs.241505.

13. Adib, R., et al. Mitotic phosphorylation by NEK6 and NEK7 reduces the microtubule affinity of EML4 to promote chromosome congression. Sci. Signal. 12, eaaw2939 (2019).

14. Papageorgiou, S., Pashley, S. L., O’Regan, L., Straatman, K. R. & Fry, A. M. Microtubule Association of EML4–ALK V3 Is Key for the Elongated Cell Morphology and Enhanced Migration Observed in V3 Cells †. Cells 13, (2024).

15. Fry, A. M., O’Regan, L., Sabir, S. R. & Bayliss, R. Cell cycle regulation by the NEK family of protein kinases. J. Cell Sci. 125, 4423–4433 (2012).

16. O’Regan, L. & Fry, A. M. The Nek6 and Nek7 protein kinases are required for robust mitotic spindle formation and cytokinesis. Mol. Cell. Biol. 29, 3975–90 (2009).

17. Belham, C. et al. A Mitotic Cascade of NIMA Family Kinases. J. Biol. Chem. 278, 34897–34909 (2003).

18. O’Regan, L. et al. Hsp72 is targeted to the mitotic spindle by Nek6 to promote K-fiber assembly and mitotic progression. J. Cell Biol. 209, 349–358 (2015).

19. Cullati, S. N., Kabeche, L., Kettenbach, A. N. & Gerber, S. A. A bifurcated signaling cascade of NIMA-related kinases controls distinct kinesins in anaphase. J. Cell Biol. 216, 2339–2354 (2017).

20. Haq, T. et al. Mechanistic basis of Nek7 activation through Nek9 binding and induced dimerization. Nat. Commun. 6, 8771 (2015).

21. Richards, M. W. et al. An Autoinhibitory Tyrosine Motif in the Cell-Cycle-Regulated Nek7 Kinase Is Released through Binding of Nek9. Mol. Cell 36, 560–570 (2009).

22. Bertran, M. T. et al. Nek9 is a Plk1-activated kinase that controls early centrosome separation through Nek6/7 and Eg5. EMBO J. 30, 2634–2647 (2011).

23. Kirkbride, K. C., Sung, B. H., Sinha, S. & Weaver, A. M. Cortactin: a multifunctional regulator of cellular invasiveness. Cell Adh. Migr. 5, 187–198.

24. MacGrath, S. M. & Koleske, A. J. Cortactin in cell migration and cancer at a glance. J. Cell Sci. 125, 1621–6 (2012).

25. Bandela, M., Belvitch, P., Garcia, J. G. N. & Dudek, S. M. Cortactin in Lung Cell Function and Disease. Int. J. Mol. Sci. 23, (2022).

26. García Ponce, A., et al. Loss of cortactin causes endothelial barrier dysfunction via disturbed adrenomedullin secretion and actomyosin contractility. Sci. Rep. 6, 29003 (2016).

27. Liu, T. et al. Cortactin stabilizes actin branches by bridging activated Arp2/3 to its nucleated actin filament. Nat. Struct. Mol. Biol. 2024 315 31, 801–809 (2024).

28. Weaver, A. M. et al. Cortactin promotes and stabilizes Arp2/3-induced actin filament network formation. Curr. Biol. 11, 370–4 (2001).

29. Clark, E. S., Whigham, A. S., Yarbrough, W. G. & Weaver, A. M. Cortactin is an essential regulator of matrix metalloproteinase secretion and extracellular matrix degradation in invadopodia. Cancer Res. 67, 4227–4235 (2007).

30. Yamada, H., Takeda, T., Michiue, H., Abe, T. & Takei, K. Actin bundling by dynamin 2 and cortactin is implicated in cell migration by stabilizing filopodia in human non-small cell lung carcinoma cells. Int. J. Oncol. 49, 877–886 (2016).

31. Kelley, L. C., Hayes, K. E., Ammer, A. G., Martin, K. H. & Weed, S. A. Cortactin Phosphorylated by ERK1/2 Localizes to Sites of Dynamic Actin Regulation and Is Required for Carcinoma Lamellipodia Persistence. PLoS One 5, e13847 (2010).

32. Bryce, N. S. et al. Cortactin Promotes Cell Motility by Enhancing Lamellipodial Persistence. Curr. Biol. 15, 1276–1285 (2005).

33. Wu, H., Reynolds, A. B., Kanner, S. B., Vines, R. R. & Parsons, J. T. Identification and characterization of a novel cytoskeleton-associated pp60src substrate. Mol. Cell. Biol. 11, 5113–5124 (1991).

34. Weed, S. A. et al. Cortactin localization to sites of actin assembly in lamellipodia requires interactions with F-actin and the Arp2/3 complex. J. Cell Biol. 151, 29–40 (2000).

35. Scherer, A. N., Anand, N. S. & Koleske, A. J. Cortactin stabilization of actin requires actin-binding repeats and linker, is disrupted by specific substitutions, and is independent of nucleotide state. J. Biol. Chem. 293, 13022–13032 (2018).

36. Martinez-Quiles, N., Henry Ho, H.-Y., Kirschner, M. W., Ramesh, N. & Geha, R. S. Erk/Src Phosphorylation of Cortactin Acts as a Switch On-Switch Off Mechanism That Controls Its Ability To Activate N-WASP. Mol. Cell. Biol. 24, 5269–5280 (2004).

37. Alexander, J. et al. Spatial Exclusivity Combined with Positive and Negative Selection of Phosphorylation Motifs Is the Basis for Context-Dependent Mitotic Signaling. Sci. Signal. 4, ra42 (2011).

38. van de Kooij, B. et al. Comprehensive substrate specificity profiling of the human nek kinome reveals unexpected signaling outputs. Elife 8, (2019).

39. Meiler, E., Nieto-Pelegrín, E. & Martinez-Quiles, N. Cortactin tyrosine phosphorylation promotes its deacetylation and inhibits cell spreading. PLoS One 7, e33662 (2012).

40. Zhang, X. et al. HDAC6 modulates cell motility by altering the acetylation level of cortactin. Mol. Cell 27, 197–213 (2007).

41. Pashley, S. L. et al. The mesenchymal morphology of cells expressing the EML4–ALK V3 oncogene is dependent on phosphorylation of Eg5 by NEK7. J. Biol. Chem. 300, (2024).

42. Garcin, C. & Straube, A. Microtubules in cell migration. Essays in Biochemistry vol. 63 509–520 (2019).

43. Dogterom, M. & Koenderink, G. H. Actin-microtubule crosstalk in cell biology. Nat. Rev. Mol. Cell Biol. 20, 38–54 (2019).

44. Schnoor, M., Stradal, T. E. & Rottner, K. Cortactin: Cell Functions of A Multifaceted Actin-Binding Protein. Trends Cell Biol. 28, 79–98 (2018).

45. Biosse Duplan, M., et al. Microtubule Dynamic Instability Controls Podosome Patterning in Osteoclasts through EB1, Cortactin, and Src. Mol. Cell. Biol. 34, 16–29 (2014).

46. Jaworski, J. et al. Dynamic Microtubules Regulate Dendritic Spine Morphology and Synaptic Plasticity. Neuron 61, 85–100 (2009).

47. Freixo, F. et al. NEK7 regulates dendrite morphogenesis in neurons via Eg5-dependent microtubule stabilization. Nat. Commun. 9, 1–17 (2018).

48. Schätzle, P. et al. Activity-Dependent Actin Remodeling at the Base of Dendritic Spines Promotes Microtubule Entry. Curr. Biol. 28, 2081--2093.e6 (2018).

49. Adams, G. et al. The microtubule plus end tracking protein TIP150 interacts with cortactin to steer directional cell migration. J. Biol. Chem. 291, 20692– 20706 (2016).

50. Inoue, H. et al. A MAP1B-cortactin-Tks5 axis regulates TNBC invasion and tumorigenesis. J. Cell Biol. 223, (2024).

51. Weed, S. A., Du, Y. & Parsons, J. T. Translocation of cortactin to the cell periphery is mediated by the small GTPase Rac1. J. Cell Sci. 111 (Pt 1, 2433–2443 (1998).

52. Grassart, A. et al. Pak1 Phosphorylation Enhances Cortactin-N-WASP Interaction in Clathrin-Caveolin-Independent Endocytosis. Traffic 11, 1079– 1091 (2010).

53. MacGrath, S. M. & Koleske, A. J. Arg/Abl2 Modulates the Affinity and Stoichiometry of Binding of Cortactin to F-Actin. Biochemistry 51, 6644–6653 (2012).

54. Wang, W., Chen, L., Ding, Y., Jin, J. & Liao, K. Centrosome separation driven by actin-microfilaments during mitosis is mediated by centrosome-associated tyrosine-phosphorylated cortactin. J. Cell Sci. 121, 1334–1343 (2008).

55. van Rossum, A. G. S. H., Schuuring-Scholtes, E., van Buuren-van Seggelen, V., Kluin, P. M. & Schuuring, E. Comparative genome analysis of cortactin and HS1: The significance of the F-actin binding repeat domain. BMC Genomics 6, 15 (2005).

56. Pant, K., Chereau, D., Hatch, V., Dominguez, R. & Lehman, W. Cortactin Binding to F-actin Revealed by Electron Microscopy and 3D Reconstruction. J. Mol. Biol. 359, 840–847 (2006).

57. Cowieson, N. P. et al. Cortactin adopts a globular conformation and bundles actin into sheets. J. Biol. Chem. 283, 16187–16193 (2008).

58. Hohmann & Dehghani. The Cytoskeleton—A Complex Interacting Meshwork. Cells 8, 362 (2019).

59. Etienne-Manneville, S. Microtubules in Cell Migration. Annu. Rev. Cell Dev. Biol. 29, 471–499 (2013).

60. Tehrani, S., Faccio, R., Chandrasekar, I., Ross, F. P. & Cooper, J. A. Cortactin has an essential and specific role in osteoclast actin assembly. Mol. Biol. Cell 17, 2882–2895 (2006).

61. Artym, V. V, Zhang, Y., Seillier-Moiseiwitsch, F., Yamada, K. M. & Mueller, S. C. Dynamic interactions of cortactin and membrane type 1 matrix metalloproteinase at invadopodia: Defining the stages of invadopodia formation and function. Cancer Res. 66, 3034–3043 (2006).

62. Hering, H. & Sheng, M. Activity-Dependent Redistribution and Essential Role of Cortactin in Dendritic Spine Morphogenesis. J. Neurosci. 23, 11759–11769 (2003).

63. Citalán-Madrid, A. F. et al. Cortactin deficiency causes increased RhoA/ROCK1-dependent actomyosin contractility, intestinal epithelial barrier dysfunction, and disproportionately severe DSS-induced colitis. Mucosal Immunol. 2017 105 10, 1237–1247 (2017).

64. Ma, L. et al. Discovery of the migrasome, an organelle mediating release of cytoplasmic contents during cell migration. Cell Res. 25, 24–38 (2015).

65. Ma, Y. et al. Isolation and characterization of extracellular vesicle-like nanoparticles derived from migrasomes. FEBS J. (2023) doi:10.1111/FEBS.16756.

66. Houtman, S. H., Rutteman, M., De Zeeuw, C. I. & French, P. J. Echinoderm microtubule-associated protein like protein 4, a member of the echinoderm microtubule-associated protein family, stabilizes microtubules. Neuroscience 144, 1373–1382 (2007).

67. Janoueix-Lerosey, I., Lopez-Delisle, L., Delattre, O. & Rohrer, H. The ALK receptor in sympathetic neuron development and neuroblastoma. Cell and Tissue Research vol. 372 325–337 (2018).

68. Gustafsson, N. et al. Fast live-cell conventional fluorophore nanoscopy with ImageJ through super-resolution radial fluctuations. Nat. Commun. 7, 1–9 (2016).

69. Schindelin, J., et al. Fiji: an open-source platform for biological-image analysis. Nat. Methods 9, 676–682 (2012).

70. Xu, T. et al. SOAX: A software for quantification of 3D biopolymer networks. Sci. Rep. 5, 1–10 (2015).

71. Tinevez, J. Y. et al. TrackMate: An open and extensible platform for single-particle tracking. Methods 115, 80–90 (2017).

