## Supplementary Information for "NEK7 phosphorylation of cortactin modulates the migratory capacity of cells expressing EML4-ALK V3"

**A**

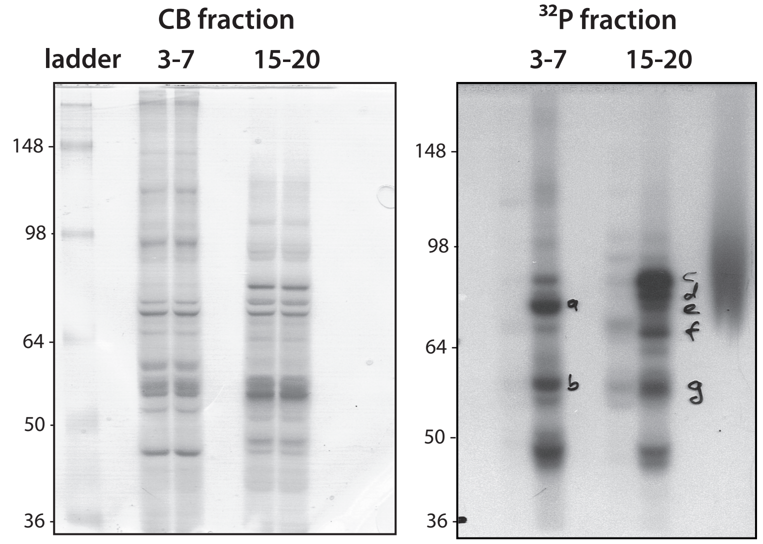


**B**

| Letter | Substrate | Initial rates units/mg |
|---|---|---|
| a | HSP70 | 2.4 |
| b | Tubulin β | 2.4 |
| c | Cortactin a | 25.3 |
| d | HSP70 | 11.3 |
| e | HSP70 | 3.3 |
| f | Cortactin a | 7.6 |
| g | Tubulin β | 3.8 |

**C**

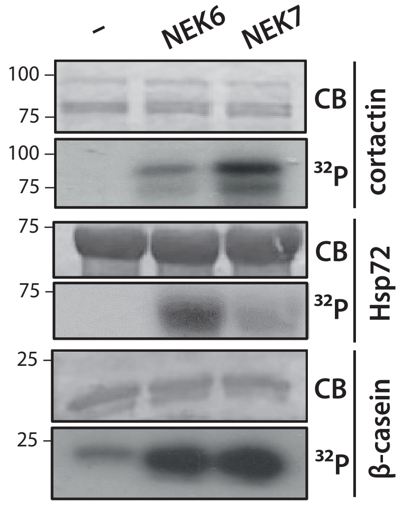


**D**

**NEK6 sites mapped**

MWKASAGHAV SIAQDDAGAD DWETDPDFVN DVSEKEQRWG AKTVQGSGHQ EHINIHKLRE NVFQEHQTLK EKELETGPKA
SHGYGGKFGV EQDRMDKSAV GHEYQSKLSK HCSQVDSVRG FGGKFGVQMD RVDQ**S**AVGFE YQGKTEKHAS QKDYSSGFGG
KYGVQADRVD K**S**AVGFDYQG KTEKHESQRD YSKGFGGKYG IDKDKVDKSA VGFEYQGKTE KHESQKDYVK GFGGKFGVQT
DRQDKCALGW DHQEKLQLHE **S**QKDYKTGFG GKFGVQ**S**ERQ DSAAVGFDYK EKLAKHESQQ DYSKGFGGKY GVQKDRMDKN
ASTFEDVTQV SSAYQKTVPV EAVTSKTSNI RANFENLAKE KEQEDRRKAE AERAQRMAKE RQEQEEARRK LEEQARAKTQ
**T**PPV**S**PAPQP TEERLP**SS**PV **Y**EDAASFKAE L**S**YRGPVSGT EPEPVYSMEA ADYREASSQQ GLAYATEAVY ESAEAPGHYP
AEDSTYDEYE NDLGITAVAL YDYQAAGDDE ISFDPDDIIT NIEMIDDGWW RGVCKGRYGL FPANYVELRQ

**E**

**NEK7 sites mapped**

MWKA**S**AGHAV **S**IAQDDAGAD DWETDPDFVN DV**S**EKEQRWG AKTVQGSGHQ EHINIHKLRE NVFQEHQTLK EKELETGPKA
SHGYGGKFGV EQDRMDK**S**AV GHEYQSKLSK HCSQVDSVRG FGGKFGVQMD RVDQ**S**AVGFE YQGKTEKHA**S** QKDYSSGFGG
KYGVQADRVD K**S**AVGFDYQG KTEKHESQRD YSKGFGGKYG IDKDKVDK**S**A VGFEYQGKTE KHESQKDYVK GFGGKFGVQT
DRQDKCALGW DHQEKLQLHE **S**QKDYKTGFG GKFGVQ**S**ERQ D**S**AAVGFDYK EKLAKHESQQ DY**S**KGFGGKY GVQKDRMDKN
A**S**TFEDV**T**QV **SS**AYQKTVPV EAV**T**SK**T**SNI RANFENLAKE KEQEDRRKAE AERAQRMAKE RQEQEEARRK LEEQARAKTQ
**T**PPV**S**PAPQP TEERLPSSPV YEDAA**S**FKAE L**S**YRGPVSGT EPEPVYSMEA ADYREASSQQ GLAYATEAVY ESAEAPGHYP
AEDSTYDEYE NDLGITAVAL YDYQAAGDDE ISFDPDDIIT NIEMIDDGWW RGVCKGRYGL FPANYVELRQ

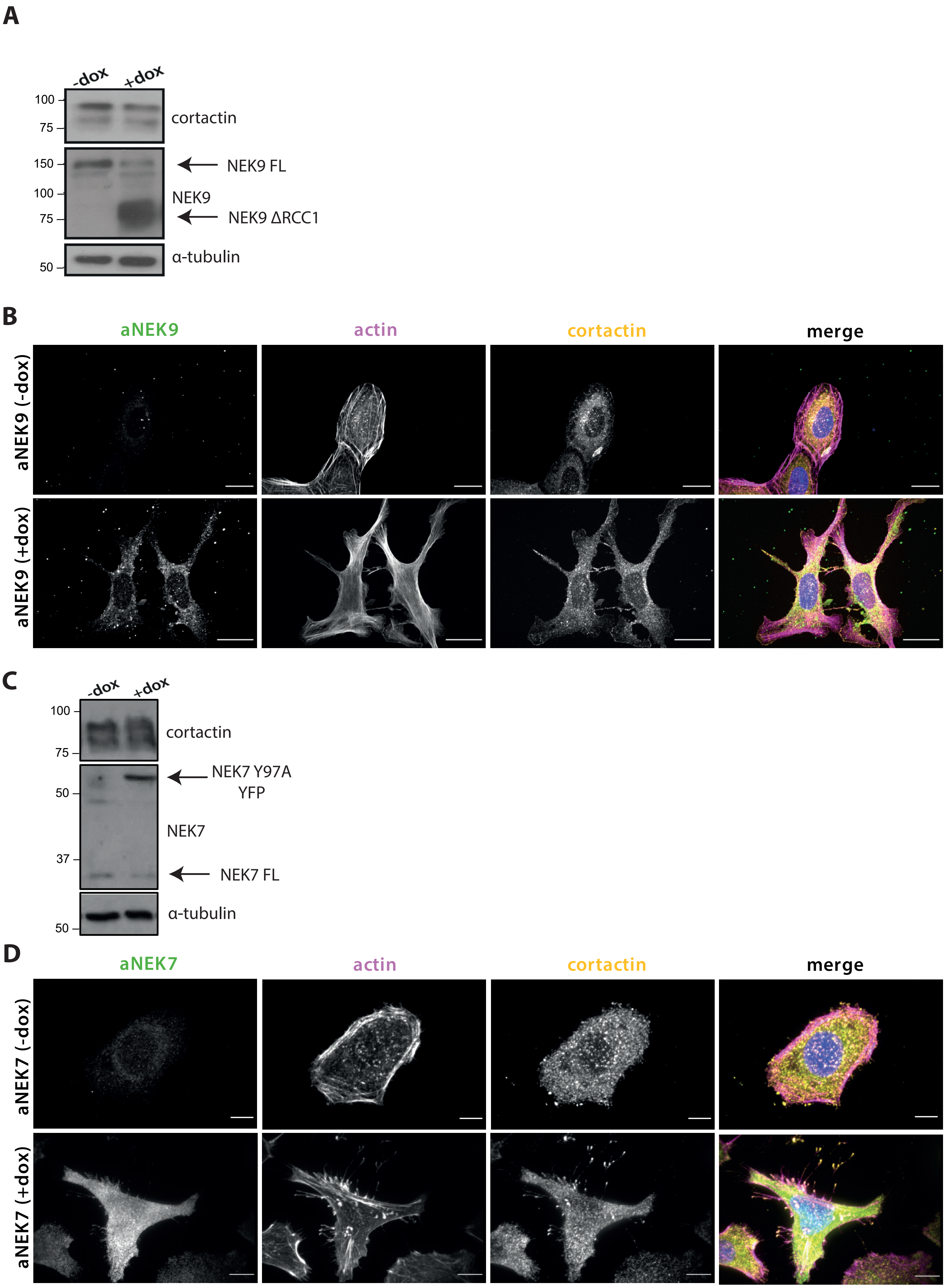
A
-dox
+dox
100
75
cortactin
150
100
75
NEK9 FL
NEK9
NEK9 ΔRCC1
50
α-tubulin
B
aNEK9
actin
cortactin
merge
aNEK9 (-dox)
aNEK9 (+dox)
C
-dox
+dox
100
75
cortactin
50
37
NEK7 Y97A
YFP
NEK7
NEK7 FL
50
α-tubulin
D
aNEK7
actin
cortactin
merge
aNEK7 (-dox)
aNEK7 (+dox)

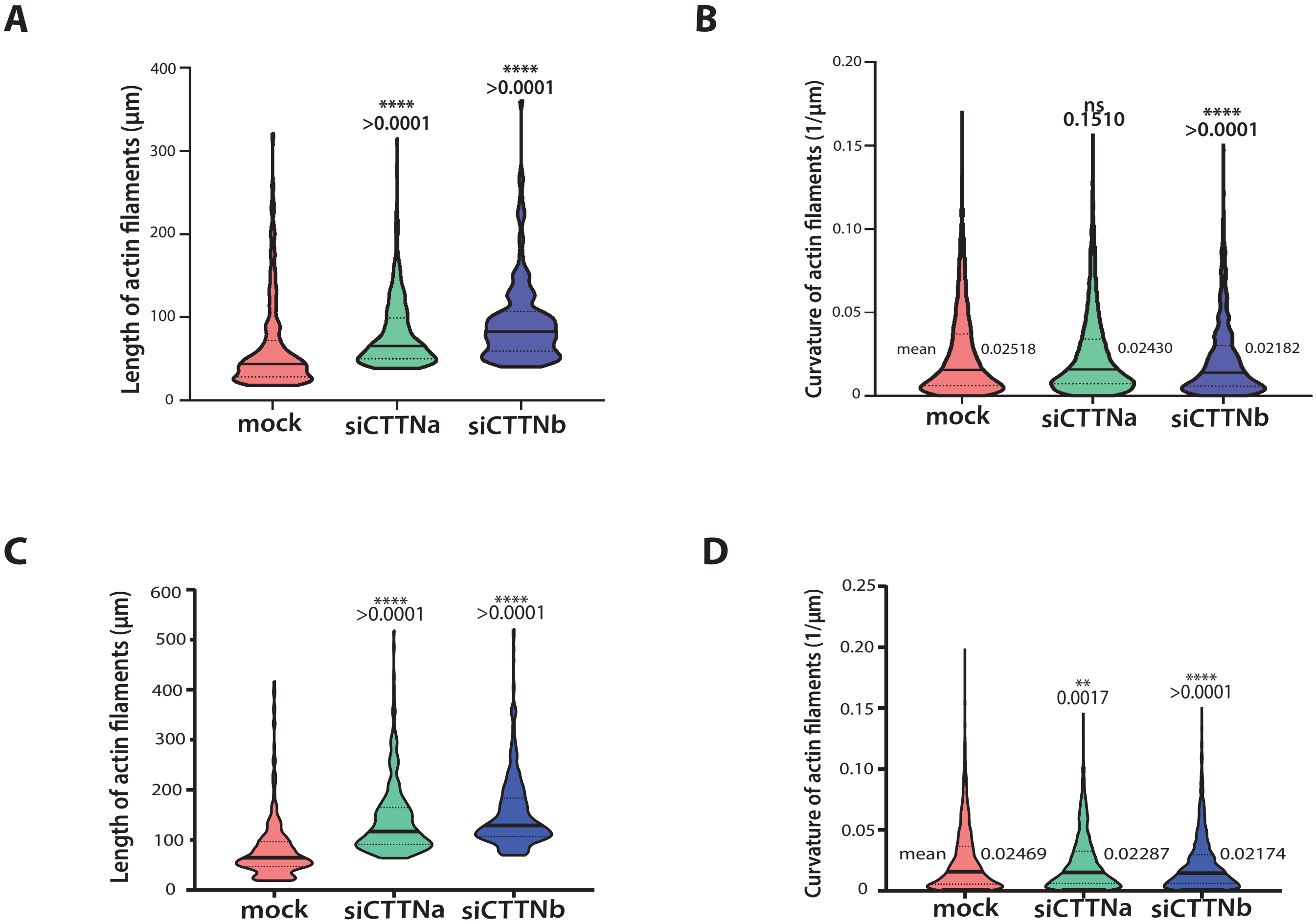
A
Length of actin filaments (µm)
400
300
200
100
0
****
>0.0001
****
>0.0001
mock
siCTTNa
siCTTNb
B
Curvature of actin filaments (1/µm)
0.20
0.15
0.10
0.05
0
ns
0.1510
****
>0.0001
mean
0.02518
0.02430
0.02182
mock
siCTTNa
siCTTNb
C
Length of actin filaments (µm)
600
500
400
300
200
100
0
****
>0.0001
****
>0.0001
mock
siCTTNa
siCTTNb
D
Curvature of actin filaments (1/µm)
0.25
0.20
0.15
0.10
0.05
0
**
0.0017
****
>0.0001
mean
0.02469
0.02287
0.02174
mock
siCTTNa
siCTTNb

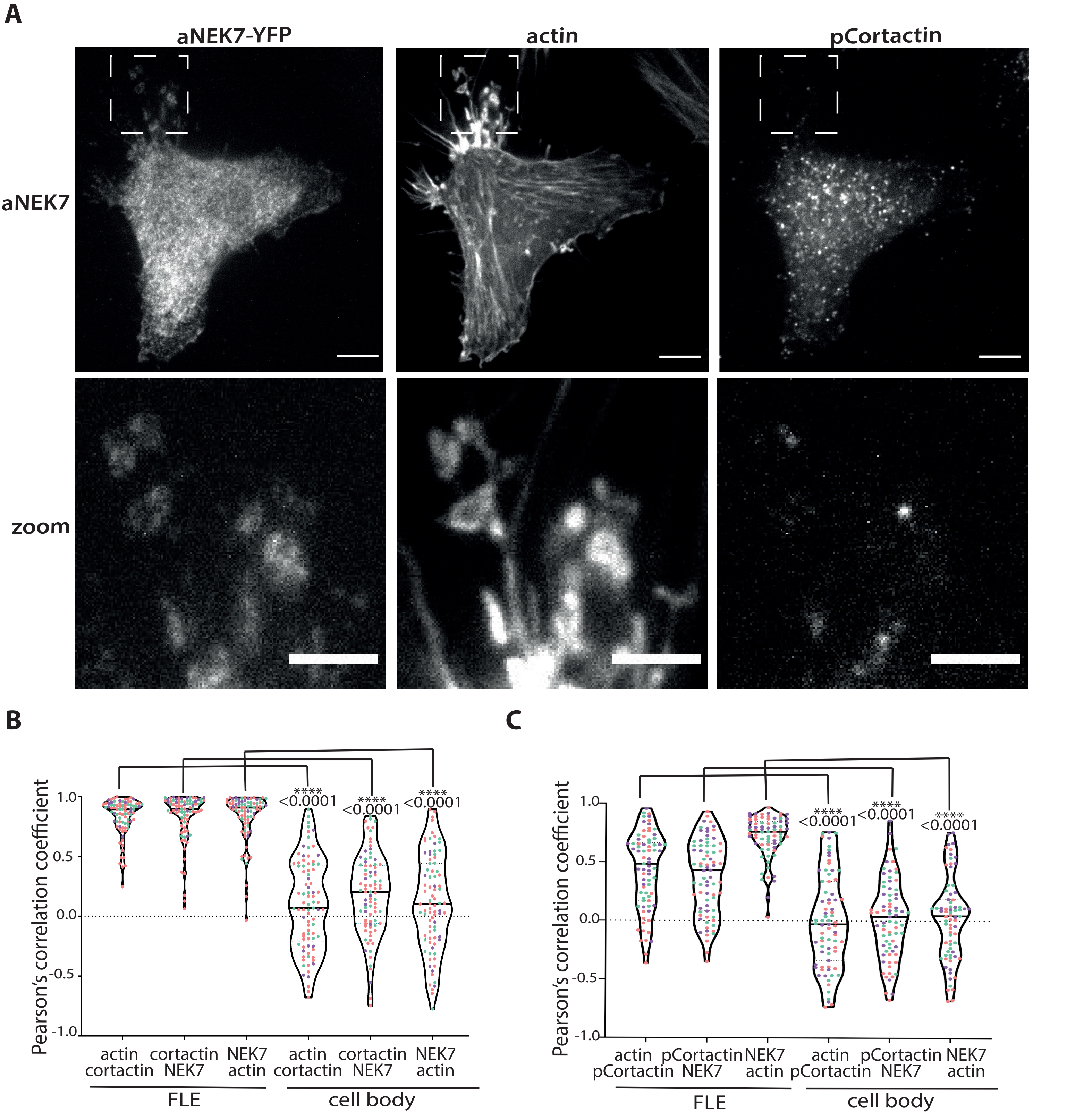
A
aNEK7-YFP
actin
pCortactin
aNEK7
zoom
B
Pearson's correlation coefficient
1.0
0.5
0.0
-0.5
-1.0
****
<0.0001
actin cortactin
cortactin NEK7
NEK7 actin
FLE
cell body
C
actin pCortactin
pCortactin NEK7
NEK7 actin

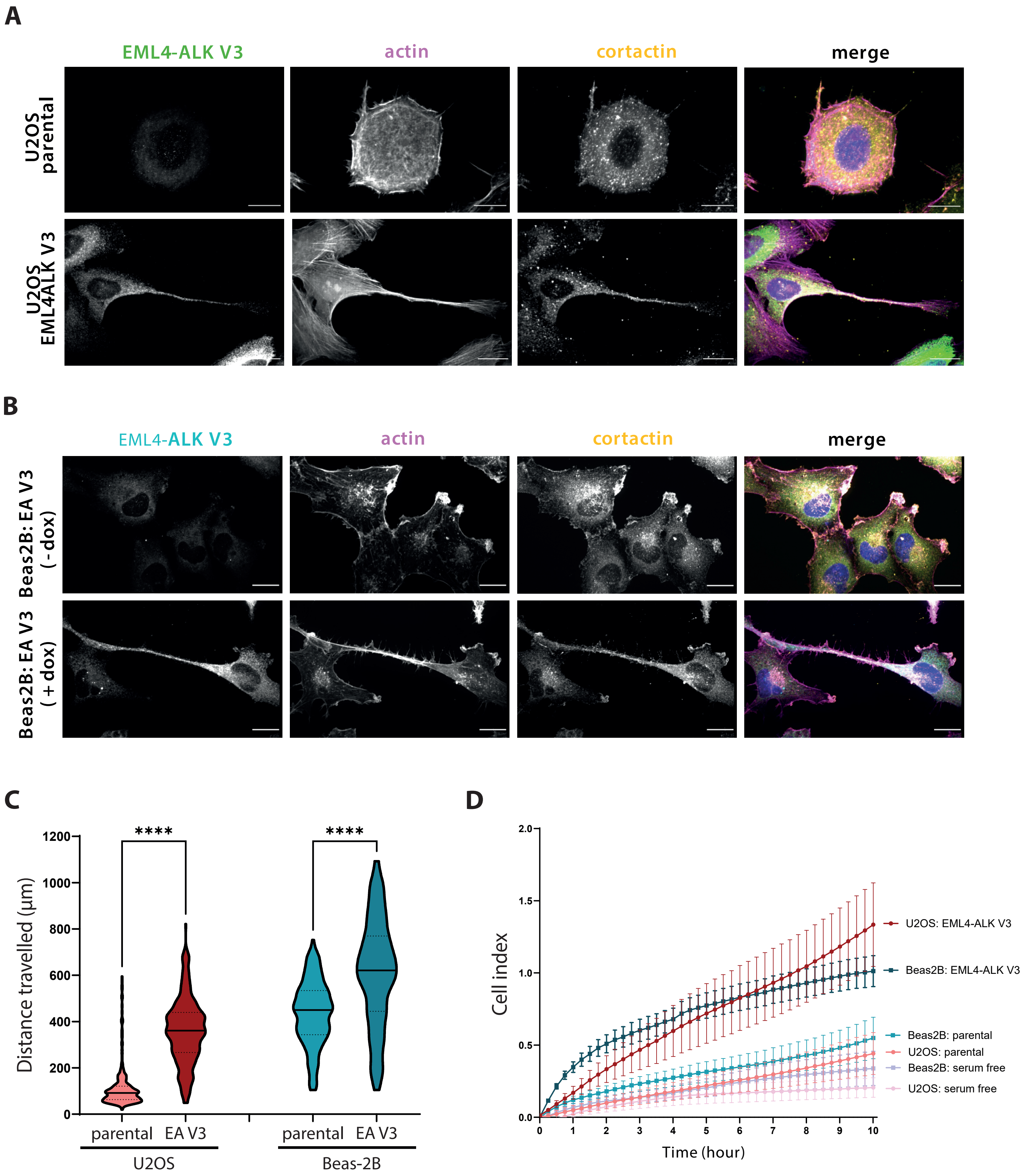
A
EML4-ALK V3
actin
cortactin
merge
U2OS parental
U2OS EML4ALK V3
B
EML4-ALK V3
actin
cortactin
merge
Beas2B: EA V3 (-dox)
Beas2B: EA V3 (+dox)
C
****
****
Distance travelled (µm)
1200
1000
800
600
400
200
0
parental
EA V3
parental
EA V3
U2OS
Beas-2B
D
Cell index
2.0
1.5
1.0
0.5
0.0
0
1
2
3
4
5
6
7
8
9
10
Time (hour)
U2OS: EML4-ALK V3
Beas2B: EML4-ALK V3
Beas2B: parental
U2OS: parental
Beas2B: serum free
U2OS: serum free

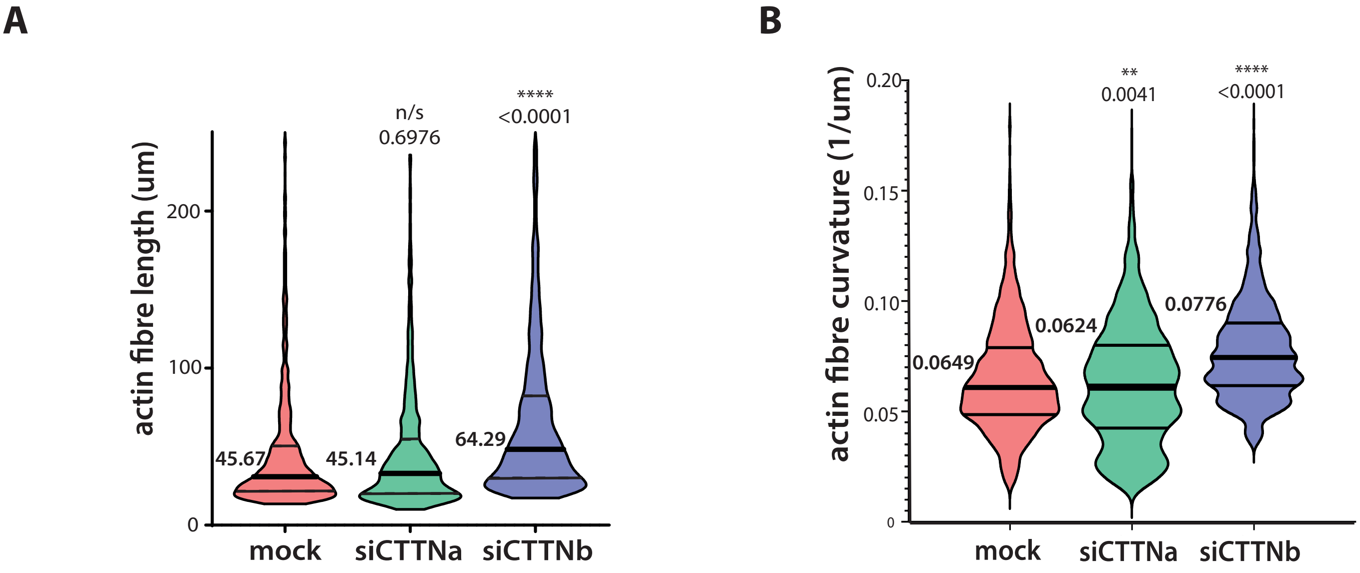
A
actin fibre length (um)
0
100
200
n/s
0.6976
****
<0.0001
45.67
45.14
64.29
mock
siCTTNa
siCTTNb
B
actin fibre curvature (1/um)
0
0.05
0.10
0.15
0.20
**
0.0041
****
<0.0001
0.0649
0.0624
0.0776
mock
siCTTNa
siCTTNb

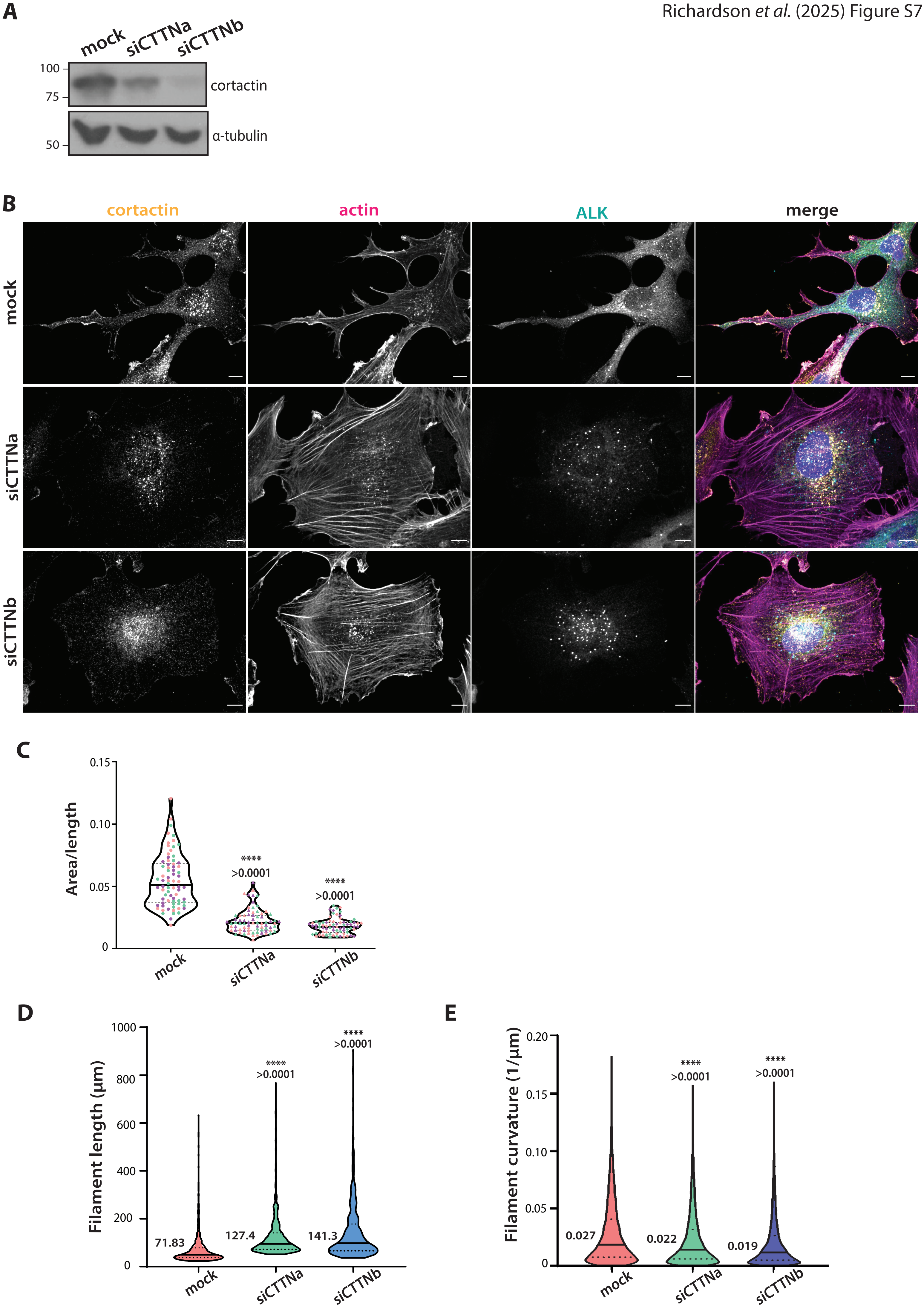
A
mock
siCTTNa
siCTTNb
100
75
50
cortactin
α-tubulin
B
cortactin
actin
ALK
merge
mock
siCTTNa
siCTTNb
C
Area/length
0.15
0.10
0.05
0
****
>0.0001
mock
siCTTNa
siCTTNb
D
Filament length (µm)
1000
800
600
400
200
71.83
127.4
141.3
E
Filament curvature (1/µm)
0.20
0.15
0.10
0.05
0.027
0.022
0.019

### SUPPLEMENTARY FIGURE LEGENDS

#### Figure S1. Identification of cortactin as a substrate of NEK6 and NEK7

**A.** HEK293 cells were lysed and fractionated as described in the Materials and Methods. Fractions from the final heparin sepharose column were incubated with purified NEK6 in the presence of $^{32}$P-γ-ATP and analysed by SDS-PAGE. Gels were stained with Coomassie Blue (CB fraction) or exposed to X-ray film ($^{32}$P fraction) to detect phosphorylated proteins. **B**. Substrates were identified using mass spectrometry fingerprinting and initial rates of phosphorylation calculated. Cortactin, HSP70 and β-tubulin were identified as the proteins indicated in panel A. **C.** Purified cortactin-WT protein was incubated with NEK6 or NEK7 proteins in vitro in the presence of [γ-32P]-ATP. Casein and Hsp72 were used as control substrates. Reactions were quenched after 1 hr with SDS-sample buffer, boiled and separated on SDS-PAGE gels for analysis by Coomassie blue stain (CB) and autoradiography ($^{32}$P). Molecular weights (kDa) are indicated on the left. **D & E.** Mass spectrometry was used to detect cortactin residues phosphorylated by NEK6 (C) and NEK7 (D) are highlighted in red. Sites phosphorylated by NEK6 were Ser135, Ser172, Ser261, Ser277, Thr401, Ser405, Ser417, Ser418, Tyr 421, and Ser432; sites phosphorylated by NEK7 were Ser5, Ser11, Ser33, Ser98, Ser135, Ser150, Ser172, Ser209, Ser261, Ser277, Ser282, Ser303, Ser322, Thr328, Ser331, Ser332, Thr344, Thr346, Thr401, Thr405, Ser426, and Ser432.

#### Figure S2. Expression of activated NEK9 or NEK7 induces a mesenchymal morphology in cancer cells.

**A.** U2OS:myc-aNEK9-ΔRCC1 cells were induced for 72 hr (+dox) and analysed by Western blot for the expression of cortactin, NEK9 full length (FL) and NEK9 ΔRCC1 (indicated with arrows) and α-tubulin. Molecular weights (kDa) are indicated on the left. **B**. U2OS:myc-aNEK9-ΔRCC1 cells were induced for 72 hr before being analysed by immunofluorescence microscopy with antibodies against aNEK9 (green) and cortactin (yellow). DNA was stained with Hoechst 33258 (blue) and actin with phalloidin-TRITC (magenta). Scale bar, 10 µm. **C** HeLa:YFP-aNEK7-Y97A were induced for 72hr (+dox) and analysed by Western blot for the expression of cortactin, NEK7 full length (FL) and Y97A-YFP (indicated by arrows). Molecular weights (kDa) are indicated on the left**. D.** HeLa:YFP-aNEK7-Y97A were induced for 72hr before being analysed by immunofluorescence microscopy with antibodies against aNEK7 (green) and cortactin (yellow). DNA was stained with Hoechst 33258 (blue) and actin with phalloidin-TRITC (magenta).Scale bar, 5 µm.

**Figure S3. Depletion of cortactin alters F-actin architecture in cells expressing activated NEK9 or NEK7**

U2OS:myc-aNEK9-ΔRCC1 (A, B) or HeLa:YFP-aNEK7-Y97A (C, D) cells were induced for 72 hr and either mock depleted or depleted of cortactin using siCTTNa or siCTTNb**. A & B.** Violin plots representing the length (A) and curvature (B) of actin filaments in U2OS:myc-aNEK9-ΔRCC1 cells. **C & D.** Violin plots representing the length (C) and curvature (D) of actin filaments in HeLa:YFP-aNEK7-Y97A cells. Solid lines represent the median and dotted lines the quartiles; mean values are indicated on B and D. **, $p$<0.01, ****, $p$<0.0001 by unpaired Student's T-test.

**Figure S4. Phosphorylated cortactin and activated NEK7 colocalise at the tips and branches of FLEs**

**A.** HeLa:YFP-NEK7-Y97A cells were induced for 72 hr before being analysed by immunofluorescence microscopy with antibodies against aNEK7 and pCortactin. Actin was stained with phalloidin-TRITC. Scale bars, 10 µm (upper panels) and 5 µm (lower panels). **B & C**. Violin plots depict the Pearson's correlation coefficient of co-localization between actin, NEK7 and cortactin (B) or pCortactin (C) in FLEs or cell body based on experiments shown in Figure 2G. Three colours represent separate experiments. Solid lines represents the median. n=3 with 10 cells and 5 colocalisation points per cell measured per experiment . Three colours represent separate experiments. Solid line represents the median. **, $p$<0.01, ****, $p$<0.0001 by unpaired Student's T-test.

**Figure S5. Expression of EML4-ALK in U2OS and Beas-2B cells induces a mesenchymal phenotype with elongated morphology and enhanced migration.**

**A.** U2OS parental and U2OS: EML4-ALK V3 cells were analysed by immunofluorescence microscopy with antibodies against ALK (green) or cortactin (yellow). DNA was stained with Hoechst 33258 (blue) and actin with phalloidin-TRITC (magenta). Scale bar, 10 µm**. B** Parental Beas-2B or Beas-2B:EML4-ALK V3 cells were induced with doxycycline for 72 hr and analysed by immunofluorescence microscopy with antibodies against ALK (teal) or cortactin (yellow). DNA was stained with Hoechst 33258 (blue) and actin with phalloidin-TRITC (magenta). Scale bars, 10 µm. **C.** U2OS and Beas-2B parental and EML4-ALK V3-expressing (EA V3) cells were plated on collagen-coated wells and distance travelled by individual cells measured over 12 hr by time-lapse microscopy. Solid lines represents the median and dotted lines the quartiles. ****, $p$<0.0001 by one-way ANOVA. **D.** U2OS and Beas-2B parental and EML4-ALK V3-expressing cells were placed in the upper chamber of a CIM-plate 16 without serum. The lower chamber was filled with media with serum as a chemo-attractant and the movement of cells across the chamber measured over 48 hr. The cell index over the 10-hr time-points is shown. Migration of mock-depleted cells with serum-free media in the lower chamber was also measured.

**Figure S6. Depletion of cortactin alters F-actin architecture in cells expressing EML4-ALK V3**

**A & B.** U2OS cells expressing EML4-ALK V3 were either mock depleted or depleted of cortactin using siCTTNa or siCTTNb as described in Figure 4**.** Violin plots represent the actin fibre lengths (A) and curvature (B). Medians are indicated with solid line and quartiles with thin solid lines. Means are indicated to the left of each plot. **p<0.01; ****, $p$<0.0001 by unpaired Student's T-test.

**Figure S7. The mesenchymal-like morphology of Beas-2B cells expressing EML4-ALK V3 is dependent on cortactin**

**A.** Beas-2B:EML4-ALK V3 cells were induced with doxycycline and either mock-depleted or depleted of cortactin using siCTTNa or siCTTNb for 72 hr. Lysates from cells were analysed by Western blot with antibodies against cortactin and α-tubulin. Molecular weight markers (kDa) are indicated on left. **B.** Cells as treated in (A) were analysed by immunofluorescence microscopy with antibodies against cortactin (yellow) and ALK (cyan); DNA was stained with Hoechst 33258 (blue) and actin with phalloidin-TRITC (magenta). Scale bars, 10 µm. **C.** Violin plot representing the mean area/length ratio of cells shown in B. Three colours represent separate experiments. **D & E.** Violin plots representing the lengths (D) and curvature (E) of actin filaments in cells shown in B. Median is indicated with solid line and quartiles with dotted lines. Means are indicated to the left of plots. $p$<0.0001 by unpaired Student's T-test.

**Supplementary Movie 1. Cells expressing activated NEK7 produce filopodia-like extensions on the cell periphery**

HeLa:YFP-aNEK7-Y97A cells were induced to express aNEK7 for 48 hr before being plated for live microscopy. Images were taken every 10 s for a 5 min period in phase (grey) and at 488 nm (green). Image depicts FLEs at the edge of a cell expressing aNEK7. Scale bar, 5 µm.

**Supplementary Movie 2. Cells expressing activated NEK7 produce filopodia-like extensions along cytoplasmic protrusions**

HeLa:YFP-aNEK7-Y97A cells were induced to express aNEK7 for 48 hr before being plated for live microscopy. Images were taken every 10 s for a 5 min period in phase (grey) and at 488 nm (green). Image depicts FLEs on the cytoplasmic protrusion of a cell expressing aNEK7. Scale bar, 5 µm.

**Supplementary Movie 3. Cells expressing cortactin S*4A form crescent shapes with long trailing ends and migrate in a random manner**

U2OS cells expressing cortactin-S4A were plated on collagen and their migration analysed. Images were taken every 30 min over a 24-hr period.

**Supplementary Movie 4. Cells expressing cortactin S*4A exhibit rapid closure of a scratch wound**

U2OS cells expressing recombinant cortactin-WT (left movie), -S*4D (middle movie) or -S*4A (right movie) were depleted of endogenous cortactin using siCTTNb oligonucleotides for 72 hr. Cells were seeded in a confluent layer before a single scratch wound was made through the centre of each well and the movement of cells measured over 24 hr.
